# Systems assessment of transcriptional regulation on central carbon metabolism by Cra and CRP

**DOI:** 10.1101/080929

**Authors:** Donghyuk Kim, Sang Woo Seo, Hojung Nam, Gabriela I. Guzman, Ye Gao, Bernhard O. Palsson

## Abstract

Two major transcriptional regulators of carbon metabolism in bacteria are Cra and CRP. CRP is considered to be the main mediator of catabolite repression. Unlike for CRP, available *in vivo* DNA binding information of Cra is scarce. Here we generate and integrate ChIP-exo and RNA-seq data to identify 39 binding sites for Cra and 97 regulon genes that are regulated by Cra in *Escherichia coli.* An integrated metabolic-regulatory network was formed by including experimentally-derived regulatory information and a genome-scale metabolic network reconstruction. Applying analysis methods of systems biology to this integrated network showed that Cra enables the optimal bacterial growth on poor carbon sources by redirecting and repressing the glycolysis flux, by activating the glyoxylate shunt pathway, and by activating the respiratory pathway. In these regulatory mechanisms, the overriding regulatory activity of Cra over CRP is fundamental. Thus, elucidation of interacting transcriptional regulation of core carbon metabolism in bacteria by two key transcription factors was possible by combining genome-wide experimental measurement and simulation with a genome-scale metabolic model.

## INTRODUCTION

Catabolite repression is a universal phenomenon, found in virtually all living organisms, ranging from the simplest bacteria to plants and animals (1,2). There is accumulating evidence to support that numerous mechanisms of catabolite repression existing within a single bacterium. A mechanism involving cyclic AMP (cAMP) and its receptor protein (CRP, cAMP receptor protein) in *Escherichia coli* was established about four decades ago (Figure 1A) (3). Since the general acceptance that cAMP-CRP provides the principal means to effect catabolite repression in *E. coli* and the closely related enteric bacteria, many aspects of CRP have been studied, including protein structure and allosteric activation (4), mechanisms of transcriptional regulation (5), and catabolite repression (6). Thus, CRP became one of the best characterized transcription factors (TF) in bacteria. The transcriptional regulator CRP is reported to regulate the expression of over 180 genes (7,8). A Chromatin Immuno-Precipitation (ChIP) method was used to determine *in vivo* binding sites of CRP in *E. coli* K-12 MG1655 (7) and other strains (9).

**Figure 1.**
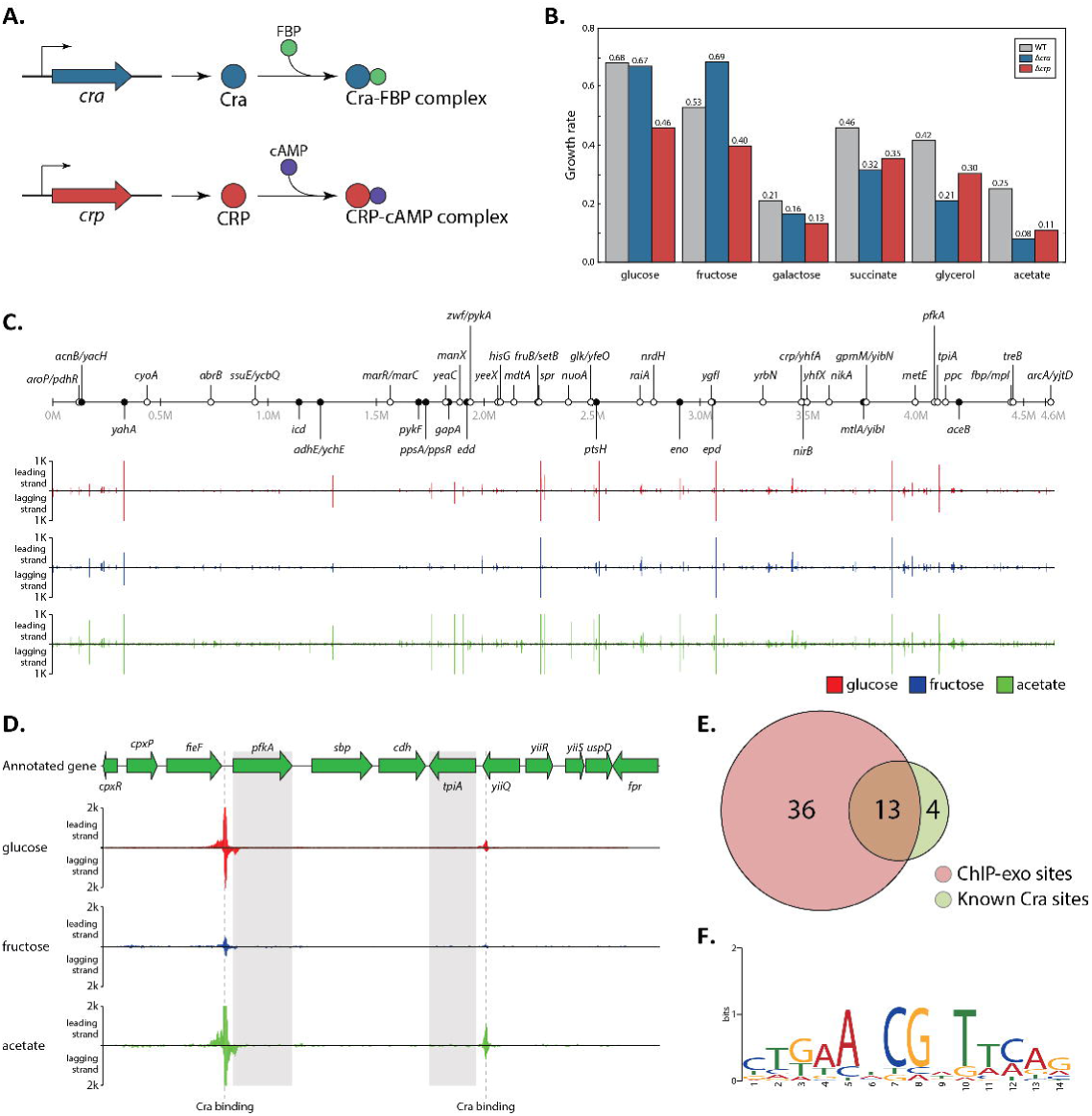
The genome-wide landscape of Cra binding in *E. coli*. (A) Cofactors of Cra and CRP. Fructose 1,6-bisphosphate (FBP) binds to Cra to deactivate Cra, while cAMP binds to CRP to activate CRP. (B) Growth rate measurement of *E. coli* WT, Δ*cra*, and Δ*crp* on different carbon sources. Disruption of *cra* showed more decrease of growth rate on less favorable carbon sources (succinate, glycerol, and acetate) than that of *crp*. (C) An overview of Cra binding profiles across the *E. coli* genome on glucose (red), fructose (blue), and acetate (green). Enrichment fold on the y-axis was calculated from ChIP-exo binding intensity in signal to noise ratio and was plotted on each location across the 4.64 Mb *E. coli* genome. Circles indicate previously identified (black) and newly identified (white) binding sites. (D) Examples of Cra binding sites upstream of *pfkA* encoding 6-phosphofructokinase-1 and *tpiA* encoding triose phosphate isomerase. In both examples, Cra binding on acetate showed the strongest intensity, while binding on fructose was the weakest. (E) Overlap between Cra binding sites from ChIP-exo experiments and previously reported sites. Out of 17 previously identified binding sites with strong evidence from the public database (29), 13 (76.5%) sites were also identified from the ChIP-exo experiments, leaving 36 binding sites newly identified. (F) From 49 Cra ChIP-exo binding sites, the sequence motif was calculated. This sequence motif is identical to one that was already known.

The carbon metabolism of enterobacteria, including *E. coli,* is globally regulated by two major TFs (1). One catabolite repression/activation mechanism identified in *E. coli* is CRP, and the other is mediated by Cra (catabolite repressor activator), which was initially named FruR (fructose repressor) (10). Cra plays a pleiotropic role to modulate the direction of carbon flow in multiple metabolic pathways, particularly in glycolysis. However, it has been postulated that Cra works independent of the CRP regulation (11,12). Multiple studies with expression profiling experiments showed that Cra is capable of regulating a large number of genes in the gluconeogenic pathway (10), TCA cycle (13), glyoxylate shunt (14), and Entner-Doudoroff (ED) pathway (12). Cra regulates glycolytic flux by sensing the concentration level of fructose-1,6-bisphosphate or fructose-1-phosphate (Figure 1A) (15).

Unlike CRP, the definition of the Cra regulon has mostly relied on transcriptome analysis or *in vitro* assays (16), and the *in vivo* identification of the Cra regulon is yet to be performed at a genome-scale. Thus, the recently developed ChIP-exo (Chromatin Immuno-Precipitation with Exonuclease treatment) (17–19) was applied to identify *in vivo* binding sites of Cra on three different carbon sources; glucose, fructose, and acetate, at the genome-scale, to enable the definition of the Cra regulon. In addition, expression profiling on different carbon sources was performed with *E. coli* wild-type and the *cra* deletion mutant to identify causal effects of the ChIP-exo identified Cra binding sites on gene expression. Using a model-based simulation, regulation of metabolic flux states by both the Cra and CRP regulons was analyzed on 38 different carbon sources including glucose, fructose, and acetate. Flux states of pathways were established with the genome-scale metabolic network model of *E. coli* (20) using flux balance analysis (FBA) (21). Integration of experimentally derived regulatory information with *in silico* calculation of flux states of core carbon metabolism revealed the transcriptional regulation by Cra of glycolysis, the TCA cycle, and the respiratory chain with emphasis on the overriding regulatory activity of Cra over CRP.

## RESULTS

In order to assess the contribution by Cra or CRP to bacterial growth on different carbon sources, *E. coli* wild-type (WT), and two knock-out strains, Δ*cra* and Δ*crp,* were grown on 6 carbon sources. Binding sites for CRP have been identified from an *in vivo* measurement with ChIP-chip and other studies (7). Thus, to identify *in vivo* Cra binding sites, *E. coli* was grown under three different carbon sources; glucose, fructose, and acetate. Glucose is a favorable carbon source for E. *coli,* and is known to cause the most severe cAMP-dependent catabolite repression (22). Cra, which is also called FruR, was first known to repress the fructose-specific *operon fruBKA* (23), thus fructose was believed to alter the activity of Cra. Acetate was chosen as a representative of less favorable carbon sources for *E*. *coli,* which is reported to relieve catabolite repression, thus changing the flux though glycolysis and altering Cra activity (15). Moreover, these carbon sources were found to make consistent optimal growth and metabolic network utilization based on *in silico* simulation of genome-scale metabolic models (24). Further model-based simulation results were used here to analyze the function of the Cra regulon in the context of reaction fluxes through energy metabolism towards gluconeogenesis. ChIP-exo and RNA-seq experiments were performed to identify Cra binding sites and expression changes of genes on these three carbon sources at the genome-scale.

### More growth defect by *cra* knock-out on the poor carbon sources

In order to assess how crucial Cra or CRP transcriptional regulation is on growth, *E. coli* WT, Δ*cra,* and Δ*crp* were grown on 6 different carbon sources to measure the growth rate and the lag-phase time (Figure 1B). Those carbon sources, glucose, fructose, galactose, succinate, glycerol, and acetate, were chosen to span various carbon sources with different numbers of carbons. On carbon sources with 6 carbon atoms, knocking out of *crp* showed much severe growth defect. However, on carbon sources with 2-, 3-, and 4-carbon atoms which would be expected to relieve catabolism repression and to induce gluconeogenesis, knocking-out the *cra* gene showed more severe growth defect (Figure 1B) and showed a much longer lag-phase time (Figure S1).

This observation confirms the involvement and importance of transcriptional regulation by Cra and CRP as shown in the previous studies. However, growth defect by disruption of *cra* more severely defected growth than did *crp*, suggesting Cra’s more important regulatory implications in adaptation to the poor carbon sources.

### Genome-wide mapping of Cra binding sites

A total of 49 Cra binding sites were identified using the ChIP-exo method during growth on three different carbon sources: glucose, fructose, and acetate (Figure 1C, Table S1). Among them, only 29 were found on fructose, indicating least activation of Cra on that carbon source. In agreement with this observation, Cra ChIP-exo peak intensity on fructose was the weakest on average among the three substrates, while peak intensity of Cra bindings on acetate was stronger than the intensity on either glucose or fructose (ranksum test p-value < 0.05, Figure S2). *E. coli* contains two phosphofructokinase (PFK) isozymes, PFK I*/pfkA* and PFK II*/pfkB,* however, over 90% of the phosphofructokinase activity is attributed to PFK I (25). Studies on *pfkA* in *E. coli* have previously identified a Cra binding site (26) upstream of a σ^70^-dependent promoter (27,28), and ChIP-exo experiments identified this binding site with a near single-base pair resolution (Figure 1D, Figure S3A). This Cra binding overlaps with the σ^70^-dependent promoter, particularly covering -35 box of this promoter, thus possibly indicating that Cra binding would repress the expression of the downstream gene, *pfkA* (Figure S3A). Similarly, Cra binding was also observed upstream of *tpiA,* which encodes triose phosphate isomerase (Figure 1D). Cra binding overlaps with the -10 and -35 boxes of the *tpiA* promoter, suggesting its repressive effect on *tpiA* expression (Figure S3B).

The genome-wide Cra binding sites were compared to binding sites summarized in a public database (29). There are 17 previously reported Cra binding sites based on experimental evidence, and 13 (76.5%) of them were identified from ChIP-exo experiments performed in this study (Figure 1E). The four missing bindings are for *cydAB, csgDEFG, hypF,* and *pck* (11,30,31). It is possible that these four binding sites were not detected in ChIP-exo experiments because they were previously identified with *in vitro* methods and/or were identified under different growth conditions such as stationary phase or anaerobic growth. One possible drawback of determining binding sites with *in vitro* methods is that they may not represent feasible *in vivo* interactions between Cra and the genomic DNA. The ChIP-exo binding sites for Cra were also compared to another dataset that was generated with the *in vitro* SELEX system (16). This study identified a total of 164 binding sites for Cra using this *in vitro* this method. Among them, only 33 binding sites overlap with the ChIP-exo binding sites, suggesting that the *in vitro* approach may have identified false-positive binding sites.

For the ChIP-exo identified binding sites, the sequence motif was calculated using the MEME suite (32). The sequence motif obtained for Cra binding sites was ctgaAtCGaTtcag (lower-case characters indicate an information content < 1 bit) (Figure 1F). This sequence motif is nearly identical to the previously reported motif gcTGAAtCGaTTCAgc (29,33).

The ChIP-exo experiments performed here on three different carbon sources provide the first genome-wide *in vivo* measurement of Cra binding sites. A total of 49 binding sites were detected, and this dataset is in good agreement with previous knowledge in terms of the genomic locations and the sequence binding motif analysis. The better resolution that the recently developed ChIP-exo method provides enabled a more precise investigation of molecular interactions between TFs and the regulatory elements, such as promoters, of the genomic DNA.

### Orchestrated regulation of carbon metabolism by Cra and CRP

The definition of the regulon for Cra necessitates integration of the Cra binding site information with transcription unit (TU) annotation. Thus, the TUs with Cra binding sites in their upstream regulatory region were chosen from the reported TU annotation (27,29). Only Cra binding sites in the regulatory regions were used in this integration, leaving out four binding sites found in the intragenic regions of y-genes, *ynfK, yegI, yejG,* and *yihP.* If the Cra binding site is located in the divergent promoter, then TUs at both sides were considered as possible Cra regulons. This integration resulted in 63 TUs with 136 genes as candidates for inclusion in the definition of the Cra regulon.

To identify TUs with expression change upon *cra* knock-out, RNA-seq experiments were performed for *E. coli* WT and Δ*cra* knock-out strains on the three carbon sources. Any TU having a gene with an expression change ≥ 2-fold (q-value ≤ 0.01) was considered as a differentially expressed TU. Out of 63 candidate TUs, 35 TUs (containing 97 genes) were differentially expressed with Cra ChIP-exo binding sites, thus 97 genes are defined as the Cra regulon (Table S2). Clusters of Orthologous Groups (COG) analysis showed that the Cra regulon has enriched functions in energy production/conversion, carbohydrate metabolism/transport, and inorganic ion transport/metabolism (Figure S4). The average number of genes in the Cra regulon TUs was 2.77, which is much larger than the average of 1.78 genes per TU for all TUs in *E. coli*(27).

Integration of Cra binding information with differential gene expression revealed the regulatory mode of Cra on its regulon TUs (Table S3, TableS4). Out of 35 regulon TUs, Cra up-regulated 16 TUs, and down-regulated 16 TUs. The remaining three TUs are up- or down-regulated depending on which of the three carbon sources was used. For instance, *glk,* which encodes the cytoplasmic glucokinase, was down-regulated on fructose, but it was up-regulated on acetate when *cra* is missing (Table S2). This result indicates a complex regulation on the expression of this enzyme, which could be true for most of the enzymes in glycolysis/gluconeogenesis and the TCA cycle, since their activity must be finely tuned based on available carbon sources.

With the Cra regulon definition and with CRP regulatory information from the public database (29), a regulatory network for Cra and CRP for core carbon metabolism was built (Figure 2A). In brief, glycolysis is more heavily regulated and always repressed by Cra. The TCA cycle, however, is more regulated and mostly activated by CRP. This regulatory logic represents a differential transcriptional regulation of glycolysis and the TCA cycle by Cra and CRP. Another interesting aspect of this reconstructed regulatory network is that only a few genes, *fbaA, gapA, pgk, epd, aceE, aceF, aceB,* and *aceA,* are co-regulated by both Cra and CRP. The two TFs regulate their overlapping target genes in an antagonizing manner. For instance, co-regulated genes in glycolysis, *fbaA, gapA, pgk,* and *aceEF,* are repressed by Cra, however they are activated by CRP. On the other hand, CRP represses the glyoxylate shunt, *aceBA,* but it is activated by Cra.

**Figure 2.**
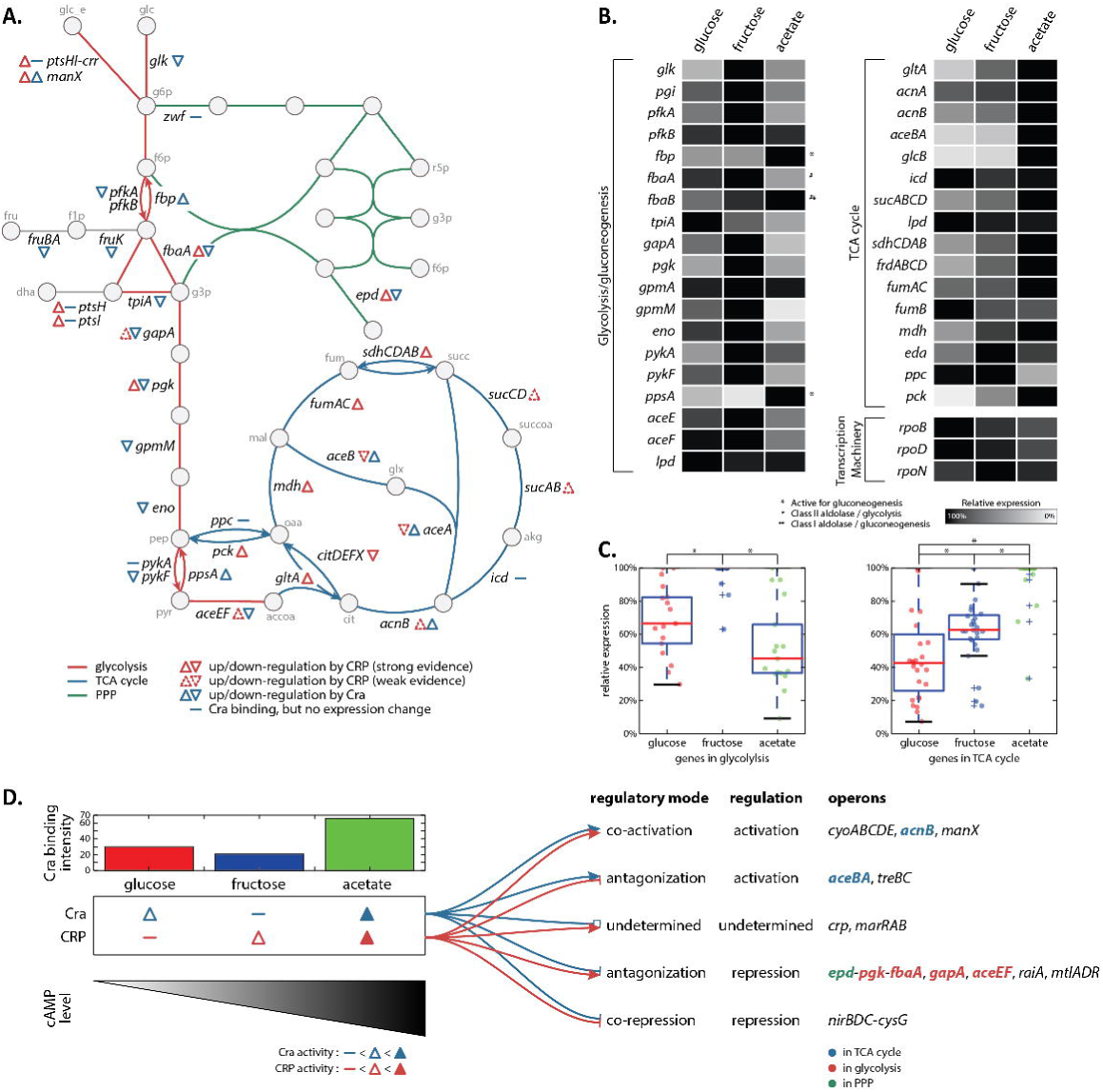
A convoluted regulation on the core carbon metabolism by Cra and CRP. (A) A metabolic network of the core carbon metabolism, glycolysis, TCA cycle, and PP pathway, with regulatory information for Cra and CRP. The information about CRP regulation was taken from the public database(29). Cra represses the glycolysis pathway, while CRP focuses on activation of the TCA cycle. Cra and CRP counteract each other in regulation of *epd-pgk-fbaA, gapA, aceEF,* and *aceBA.* (B) The relative expression of genes in glycolysis and the TCA cycle on glucose, fructose, and acetate was compared. As a control, three genes in transcription machinery, *rpoB, rpoD,* and *rpoN,* were compared in their transcriptional level. (C) Genes in glycolysis were more expressed on fructose, and were less expressed on acetate, except for a few genes that are necessary for gluconeogenesis. However, genes in the TCA cycle were the most expressed on acetate, and were the least expressed on glucose. * indicates ranksum test p-value < 0.01. (D) Depending on the regulatory mode of Cra and CRP on the shared target genes, regulatory modes are categorized into co-activation, antagonization, undetermined, and co-repression. For antagonization cases, the regulation result always follows the regulation by Cra. For instance, *epd-pgk-fbaA* operon is repressed by Cra and activated by CRP on acetate; however, the expression of this operon is down-regulated on acetate.

There is differential, but overlapping, transcriptional regulation of core carbon metabolic pathways, glycolysis, and the TCA cycle, by Cra and CRP. To investigate this complicated transcriptional regulation, expression of genes in carbon metabolism was analyzed, because gene expression is the product of the transcriptional regulation. Thus, the relative transcription of each regulated gene was compared on three different carbon sources (Figure 2B, Table S5). Except for 3 genes (*fbp*, *fbaB*, and *ppsA)* that are known to be active for gluconeogenesis, genes in glycolysis are transcribed more on fructose than glucose or acetate. *fbp* encoding fructose-1,6-bisphosphatase and *ppsA* encoding phosphoenolpyruvate synthetase catalyze the two irreversible reactions that distinguish glycolysis and gluconeogenesis. *fbaB* encodes a class I fructose bisphosphate aldolase, that is involved in gluconeogenesis, whereas class II aldolase, which is encoded by *fbaA,* is involved in glycolysis (34). Thus these genes are expected to be more highly transcribed on acetate where gluconeogenesis is active, as shown by the expression profiling data.

Acetate is primarily metabolized through the TCA cycle, to generate energy and biosynthetic precursors. Some of the acetate has to be metabolized through gluconeogenesis to synthesize five and six carbon biosynthetic precursors. Consistent with this expectation, the majority of genes in the TCA cycle were more highly expressed on acetate than on fructose or glucose. The statistical analysis of relative expression of genes in glycolysis and the TCA cycle on three carbon sources confirms the expected transcription pattern (Figure 2C). The average relative transcriptional level of glycolysis is the highest on fructose, followed by glucose, and then acetate. In this analysis, genes that are more active for gluconeogenesis were excluded for the clarity of the figure; however, those genes do not change the pattern in the relative transcriptional level even if genes in gluconeogenesis were included (Figure S5). For the genes in the TCA cycle, however, the average relative transcriptional level is the highest on acetate, followed by fructose and glucose.

The integration of ChIP-exo binding site information for Cra with known TU annotations and expression profiling by RNA-seq revealed the genome-wide transcriptional regulation by Cra. The Cra regulatory information was then combined with CRP regulatory information to build a regulatory network of the core carbon metabolic pathways including glycolysis, the TCA cycle, and PP (pentose phosphate) pathway. This regulatory network represents a differential, but overlapping, regulation of the carbon metabolism by Cra and CRP. These observations suggest a possible decoupling in regulation on glycolysis and the TCA cycle, however how this decoupling occurs in the context of the function of the entire metabolic network requires more elaboration.

### Antagonizing regulatory mode between Cra and CRP on the key enzymes of core carbon metabolism

The activity level of Cra and CRP vary depending on the carbon source. ChIP-exo experiments show that the binding activity of Cra is the lowest on fructose in terms of number of binding sites and the binding intensity, and it is strongest on acetate. However, the activity of CRP may be different. Interestingly, the expressed mRNA and protein level of both *crp* or *cra* does not change significantly during growth on glucose, fructose, or acetate (Figure S6, Figure S7). Therefore, the regulatory activity of CRP could be strongly dependent on the concentration of its effector molecule, cAMP. The intracellular concentration of cAMP is lowest on glucose, and higher on fructose. Further, the cAMP concentration is even higher on less favorable carbon sources, such as malate (22). Thus, the DNA binding and the regulatory activity of CRP is expected to be the weakest on glucose and the strongest on acetate (Figure 2D).

Cra and CRP co-regulate a total of 13 TUs. Of these, four are either co-activated or co-repressed by both of them, thus there is no conflict in regulation of those TUs between Cra and CRP. Cra binds upstream of *crp* (Figure S8), and *marRAB,* and these two TUs are reported to be activated by CRP (7). However expression of *crp* or *marRAB* did not change significantly on different carbon sources, thus they are categorized as undetermined.

Cra and CRP both regulate seven TUs containing multiple important enzymes in carbon metabolism in an opposite, or antagonizing, manner. For instance, Cra activates the *aceBA* operon that encodes enzymes in the glyoxylate shunt, while CRP represses it. On the other hand, Cra represses *epd-pgk-fbaA, gapA,* and *aceEF,* most of which are involved in glycolysis, but CRP activates their expression. This conflicting regulation by Cra and CRP makes sense on glucose and fructose, where either one of them is inactivated. Cra is more active on glucose, while CRP is more active on fructose. This differential activation explains why genes in glycolysis and the TCA cycle are more highly transcribed on fructose than on glucose.

Whereas either Cra or CRP is inactivated on glucose or fructose, acetate and possibly poor carbon sources would activate Cra and CRP at the same time. Since 7 TUs are regulated by both Cra and CRP, the expression change of genes in those TUs on less favorable carbon sources is of interest. The mRNA expression of *aceBA* and *treBC* were up-regulated on acetate, and the expression of *epd-pgk-fbaA, gapA, aceEF, raiA,* and *mtlADR* were down-regulated. Interestingly, the expression changes of these TUs always follow the regulatory mode of Cra regardless of CRP regulation. For example, expression of *aceBA* was up-regulated on acetate although CRP represses its expression, thus resulting in activation of the glyoxylate shunt. Similarly, expression of *epd-pgk-fbaA, gapA,* and *aceEF* was repressed even though CRP up-regulates their transcription, contributing to down-regulation of genes in glycolysis on acetate.

Collectively, Cra and CRP are the most active on poor carbon sources. When they are active, they regulate target genes in the core carbon metabolism in a variety of modes. They co-activate or co-repress some of their target genes, while regulating some key genes in glycolysis and the TCA cycle in an antagonizing manner. The overall regulatory consequence always follows the regulatory mode of Cra, indicating the possible overriding regulatory effect by Cra over CRP for those key enzymes of the core carbon metabolism.

### Flux balance analysis leads to a network level understanding of the regulatory roles of Cra and CRP

In order to try to understand the regulatory decoupling of glycolysis and the TCA cycle activation and to understand the metabolic driving force of increased TCA cycle activation and how the overriding regulatory effect by Cra works in that context, we turned to the methods of systems biology. Flux Balance Analysis (FBA) (21) and Markov Chain Monte Carlo (MCMC) sampling were applied to the *E. coli* metabolic model iJO1366 (20) to simulate feasible flux states for all metabolic reactions during growth on different carbon sources (Figure 3A, detailed procedures in MATERIALS AND METHODS). To determine if the predicted flux calculation correlates with the enzyme abundance and is in agreement with previous studies (24,35), the calculated flux values through the metabolic reactions were compared with the transcriptional level of the genes that encode enzymes catalyzing these reactions (Figure 3B). The ratio of fluxes through those reactions showed a good correlation to the expression change calculated between acetate and glucose. In other words, as observed in expression profiling, fluxes through the TCA cycle were predicted to increase on acetate, while fluxes through glycolysis were calculated to decrease. Enzymatic reactions in glycolysis showed reduced expression and reduced flux on acetate while reactions in the TCA cycle showed increased expression and increased flux on less favorable carbon sources. Separation between reactions in glycolysis and the TCA cycle in expression and flux changes on carbon source shift reflects a decoupling between glycolysis and the TCA cycle, and differential transcriptional regulations on them. Cra redirects the fluxes through the glycolysis pathway towards gluconeogenesis and represses the transcriptional expression of the enzymes in glycolysis to make a reduced volume of enzymatic fluxes, whereas CRP activates transcription for the majority of components in the TCA cycle resulting in more reaction flux.

**Figure 3.**
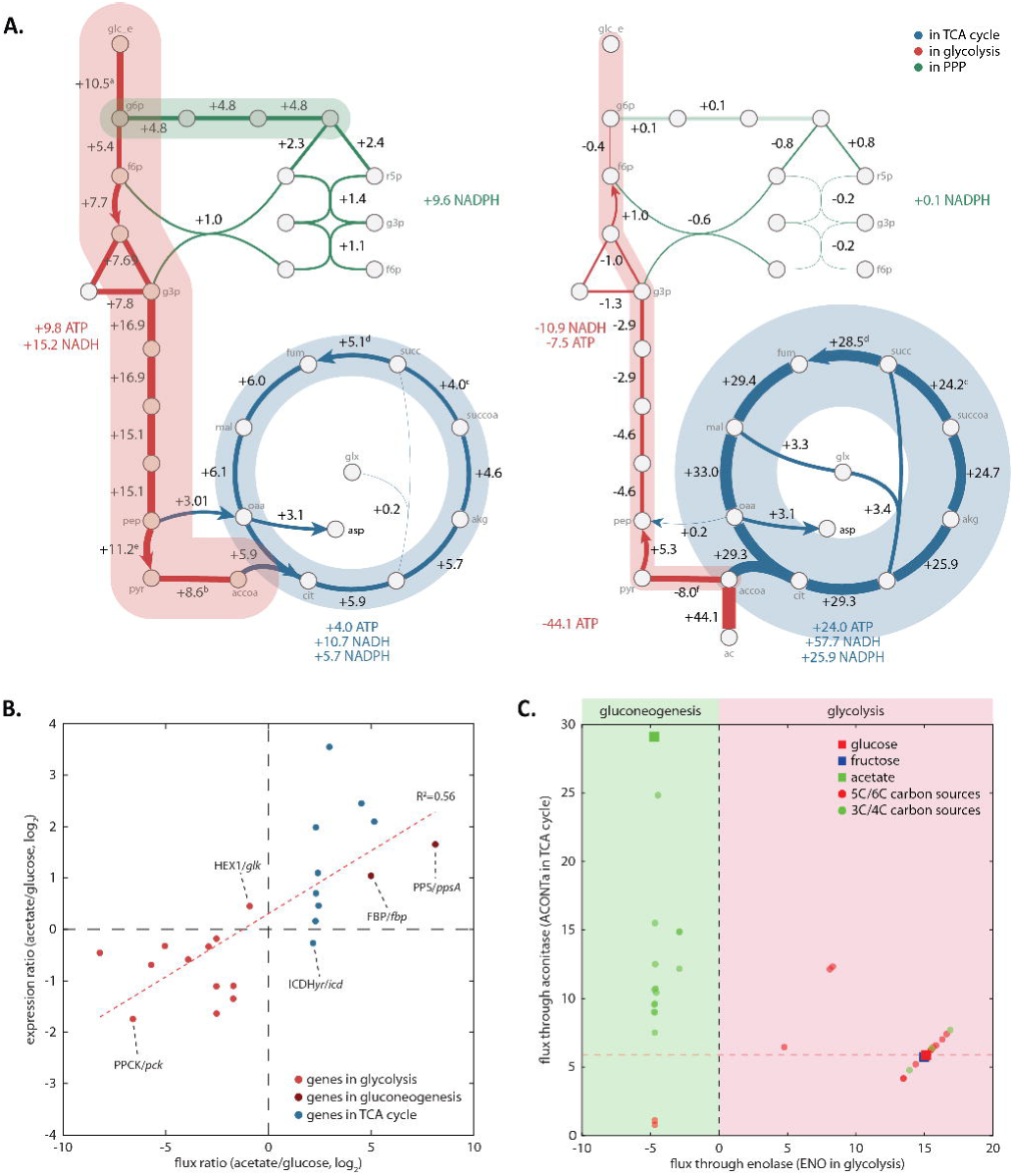
Simulated flux through the carbon metabolism explains decoupling of glycolysis and the TCA cycle and more flux through TCA cycle on poor carbon sources. (A) Normalized fluxes through glycolysis, TCA cycle, and PP pathway were calculated on glucose and acetate with a net energy production or consumption from each pathway. Consistent with the expression profiling, the TCA cycle had more activated flux on acetate than on glucose. a: HEX1+GLC*ptspp*, b: PDFI-PFL, c: -PPCSCT-SUCOAS, d: SUCD*i*-FRD2-FRD3, e: PYK+GLC*pfspp*, f: PDH-PFL+POR5, g: ACS-PTA*r* (B) Comparison between the simulated flux ratio (acetate/glucose) through the reactions in glycolysis and the TCA cycle and the expression change of genes that are responsible for those reactions. The flux ratio and expression ratio have a good correlation (R^2^ value = 0.56). Only two reactions, HEX1 and ICDFI*yr*, disagreed. Reactions in the TCA cycle are more activated in terms of flux and transcription, and reactions in glycolysis are repressed, illustrating a decoupling between reactions in glycolysis and the TCA cycle. (C) Normalized fluxes were calculated for 38 carbon sources: 1 for 2-carbon carbon source, acetate, 9 for 3-carbon carbon sources, 8 for 4-carbon carbon sources, 6 for 5-carbon carbon sources, and 12 for 6-carbon carbon sources. Enolase (ENO) and aconitase (ACONTa) were chosen as representative reactions for glycolysis (x-axis) and the TCA cycle (y-axis). 5- and 6-carbon carbon sources tend to render higher flux through glycolysis than the TCA cycle, whereas 2-, 3-, and 4-carbon carbon sources showed more flux through the TCA cycle, with much less flux through glycolysis.

Normalized fluxes, through the reactions in core carbon metabolic pathways, were mapped onto the metabolic network for glucose and acetate to illustrate how differently each pathway is predicted to be active on different carbon sources (Figure 3A). When the flux of 10.5 (mmol/gDW/hr) enters into glucose 6-phosphate (g6p) the simulation predicted 4.8 (45.7%) of the influx would flow into the PP pathway, leaving 5.4 (51.4%) to glycolysis. The simulation predicted the flux of 15.1 would flow to phosphoenolpyruvate (pep), and flux would be divided into two flux flows from pep into the TCA cycle, making flux of 5.9 from citrate to isocitrate. Thus, on glucose, glycolysis is calculated to have a higher flux than the TCA cycle. On acetate, the carbon flow starts from acetate with incoming flux of 44.1, with 8.0 (18.1%) predicted to enter into gluconeogenesis and most of the remaining flux, 29.3 (66.4%), was predicted to flow into the TCA cycle. Thus, the TCA cycle flux computed for growth on acetate is almost 5 times larger than the flux on glucose (Figure S9). There is a reaction, CITL (citrate lyase), converting citrate to oxaloacetate (oaa), however almost zero flux was predicted for this reaction in accordance with the knowledge that *E. coli* K-12 MG1655 does not have citrate lyase activity. The transcriptional level of *citCDEFXGT* for this reaction was very low (Figure S10). Thus, this reaction was ignored in further analysis.

Differential activation of the glycolysis pathway and the TCA cycle, which are regulated by Cra and CRP at the transcriptional level, was observed on three representative carbon sources, glucose, fructose, and acetate, for which experimental measurements were performed. However, *in silico* analysis with a genome-scale metabolic model on 38 carbon sources that support *E. coli* growth resulted in confirmation of the previous observation and expanded the understanding that decoupling of the glycolysis pathway and TCA cycle, reduced activity of the glycolysis pathway, and increased activity of the TCA cycle is expected to happen on most of the poor carbon sources (Figure 3C). Except for a small number of carbon sources with 3- or 4-carbons that need to be converted and fed into the glycolysis pathway, the majority of viable carbon sources with 3 or 4 carbons were predicted to render a smaller volume of fluxes through glycolysis with the opposite direction towards gluconeogenesis and to have more reaction fluxes through the TCA cycle. Thus, acetate is the best representative of the poor carbon sources that can render reduced reaction flux through glycolysis and activated flux through the TCA cycle. However, differential regulation and activation of those two pathways is not a phenomenon that is limited to a certain carbon source, acetate, but it is an outcome of a complex regulation that is common in most of the poor carbon sources.

In summary, *in silico* simulation in the context of orchestrated transcriptional regulation by Cra and CRP on the core carbon metabolism confirmed the decoupling of glycolysis and the TCA cycle at the transcriptional regulation level and reaction flux level. Repression of glycolysis and activation of the TCA cycle at the reaction flux level is happening on most of the poor carbon sources, which are mediated by Cra and CRP.

### The overriding regulatory activity of Cra over CRP on glycolysis

On the poor carbon sources such as acetate, both Cra and CRP actively try to regulate the expression of the key enzymes in glycolysis but in opposite ways. Cra tries to down-regulate the expression of the majority of enzymes in the glycolysis pathway except for two genes, *fbp* and *ppsA,* that are responsible for the flux redirection towards gluconeogenesis and should be up-regulated on the poor carbon sources to supply 5- or 6-carbon precursor molecules. CRP tries to up-regulate the expression of some metabolic enzymes in glycolysis such as *fbaA, gapA,* and *pgk.* Despite the transcriptional regulation by Cra and CRP in opposite directions, the transcriptional expression of genes for glycolysis enzymes was down-regulated following the regulatory mode of Cra (Figure 4A). The *in silico* simulation with the genome-scale metabolic model, which is independent of the transcriptional regulatory information, suggested that the reaction fluxes through the glycolysis pathway should be decreased to support the optimal growth of a bacterial cell (Figure 4B), indicating the regulatory mode by Cra is optimal while that by CRP is not.

**Figure 4.**
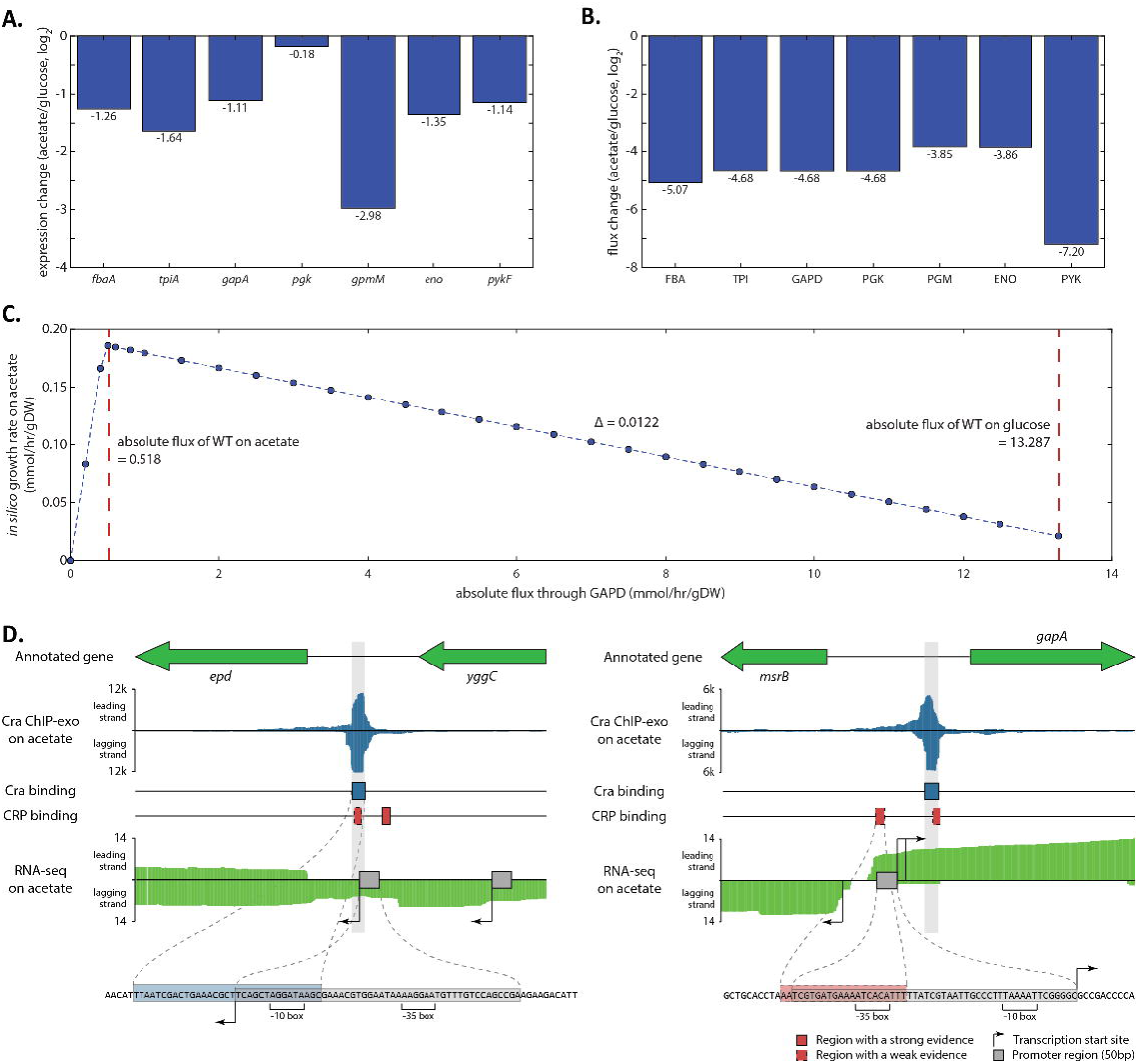
Activity of Cra overrides activity of CRP on the glycolysis pathway. (A) Expression of genes in the glycolysis pathway was repressed on acetate compared to glucose. Among the listed genes of glycolysis, *fbaA, gapA,* and *pgk* are repressed by Cra, while CRP tries to activate them. (B) Normalized reaction fluxes through glycolysis were calculated to decrease on acetate compared to glucose, agreeing with expression changes. (C) When the model was used to simulate growth on acetate, *in silico* growth rate was predicted to decrease as the glyceraldehyde 3-phosphate dehydrogenase reaction (GAPD) in glycolysis was forced to have a higher volume of reaction flux. The left red dot line indicates the reaction flux on acetate, and the right one indicates the reaction flux on glucose. (D) In-depth mapping of Cra binding sites and promoters explains how the activity of Cra can override the activity of CRP. Cra binds onto the promoter of *epd-pgk-fbaA* operon covering the transcription start site, which can interfere with transcriptional activation by CRP. Similarly, Cra binds downstream of the *gapA* promoter blocking the proceeding of RNA polymerase.

In order to verify that the reduced fluxes through glycolysis support the optimal growth, the genome-scale metabolic model was artificially forced to have a higher flux than optimal, and we computed how increased fluxes through the glycolysis pathway towards gluconeogenesis affected cell growth (Figure 4C). As postulated, when the *in silico* model was simulated on acetate but with a higher flux volume though glycolysis, the model predicted that the cell growth rate would decrease as the glycolysis flux volume increased. This indicates that the enzymatic activity of glycolysis should be lowered to support the maximum cell growth, and Cra provides the necessary transcriptional regulation on those enzymes. Thus, without the overriding activity of Cra over CRP at the transcriptional level, a bacterial cell would not be able to acquire the ability to adapt to the poor carbon sources and the optimal growth capability.

These independent lines of evidence support the notion that transcriptional regulation of those enzymes should follow the regulatory mode of Cra, ignoring the regulatory activity of CRP, and emphasize the importance of the overriding regulatory activity of Cra on glycolysis. The remaining question is, what is the molecular basis for Cra overriding activity of CRP under the condition where they are both active? With the high resolution that ChIP-exo provides, interaction between promoters (28), CRP binding sites (29), and Cra binding sites were analyzed at a base-pair resolution (Figure 4D). For *epd,* which is activated by CRP and repressed by Cra, CRP binds upstream of the core promoter, indicating this interaction between CRP and RNA polymerase (RNAP) machinery is Class I (5). Interestingly, however, Cra binding overlaps the promoter region with covering -10 box and transcription start site (TSS). This suggests Cra could stymie RNAP binding to the promoter or to initiate the transcription process even if CRP tries to recruit RNAP towards the promoter, repressing the expression of the downstream gene. Similarly, expression of *pdhR-aceEF-lpd* is repressed by Cra while being activated by CRP. CRP binds onto the genomic region including -35 box (Class II activation) (5), however Cra binds downstream of two promoters of the TU, obstructing the transcription (Figure 4D). The same regulatory interaction between CRP and Cra was observed for *mtlA* and *gapA.* Thus, the activity of Cra overrides the activity of CRP when CRP activates and Cra represses the target gene by Cra binding -10 box of the promoter, TSS, or upstream of the promoter.

### The overriding regulatory activity of Cra over CRP on the glyoxylate shunt

Expression profiling showed that the transcriptional expression of enzymes in the TCA cycle pathway was up-regulated on acetate, a poor carbon source. Computation with the genome-scale metabolic model provided support for an increased flux through the TCA cycle, leading to an increase in the cell growth rate on acetate and other poor carbon sources. Activation of the TCA cycle may require activation of the glyoxylate shunt pathway, which is encoded by *aceBA,* because that particular pathway contributes to replenishing the oxaloacetate pool.

In order to investigate this possibility, the expression profiling data was analyzed to confirm the up-regulation of transcriptional expression of the operon *aceBA* (Figure 5A). As postulated, the expression of *aceBA* was up-regulated, and Cra activates the expression of this operon while CRP tried to do the opposite. The flux prediction from *in silico* simulation confirmed that the fluxes through the glyoxylate shunt were predicted to increase when grown on acetate (Figure 5B). As an independent verification of the necessity of the glyoxylate shunt activation, the metabolic model was artificially forced to have a lower flux. The model predicted that the *in silico* growth rate would decrease as the glyoxylate shunt was forced to have a lower flux (Figure 5C).

**Figure 5.**
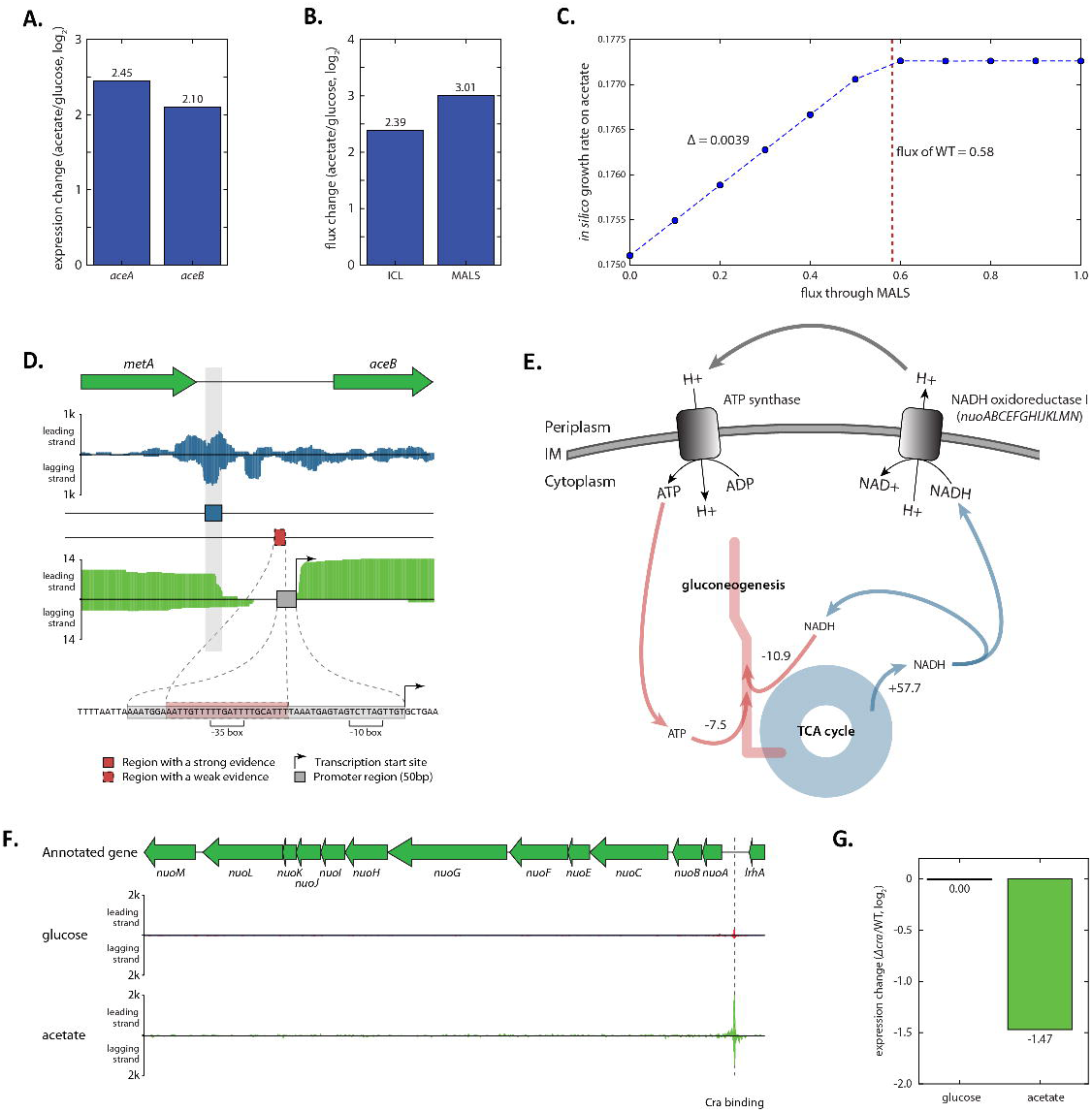
Regulatory activity of Cra on the glyoxylate shunt and the respiratory chain. (A) Expression of genes in the glyoxylate shunt, *aceA* and *aceB,* was activated on acetate compared to glucose. (B) Normalized reaction fluxes through the glyoxylate shunt were calculated to increase on acetate compared to glucose, agreeing with expression changes. (C) When the model was used to simulate growth on acetate, *in silico* growth rate was predicted to decrease as reactions of the glyoxylate shunt were forced to have a lower volume of reaction flux. (D) In-depth mapping of a Cra binding site and the promoter of aceBA explains how the activity of Cra can override the activity of CRP. (E) The model-based simulation predicted that the majority of NADH produced from the TCA cycle on glucose was fed into NADH oxidoreductase I reaction to pump out protons into the periplasm, which was used to make ATP from ATP synthase reaction. In turn, ATP is consumed in gluconeogenesis. (F) The ChIP-exo experiments detected Cra binding upstream of *nuoABCEFGHIJKLMN,* indicating regulation by Cra. (G) The expression of *nuoABCEFGHIJKLMN* operon did not change on glucose when *cra* was knocked out, however the expression was down-regulated on acetate. This suggests Cra activates the expression of the *nuoABCEFGHIJKLMN* operon.

Thus, activation of the glyoxylate shunt is required to support optimal cell growth, and the transcriptional regulation by Cra provides up-regulation of enzymes in that pathway while CRP tries to down-regulate the expression of those enzymes. Thus, the overriding regulatory activity of Cra over CRP on the glyoxylate shunt pathway is fundamental. The promoter regions and their neighboring regulatory regions of *aceBA* were also analyzed to gain insights into the molecular mechanisms of this regulatory overriding (Figure 5D). In the previous study, CRP was claimed to bind on the *aceBA* promoter covering - 35 box and to repress the expression of the target gene (36). From the ChIP-exo dataset, Cra binds upstream of the promoter, and up-regulates the expression. However, repression by CRP binding does not quash activation of *aceBA* by Cra on acetate, and this could be because CRP repression was observed to take place when *fur* is missing (36). Thus, the activity of Cra may prevail over the activity of CRP on regulation of *aceBA.*

### Cra regulates the respiratory chain to keep energy balance between reduced glycolysis and activated TCA cycle

Usage of gluconeogenesis requires energy. To make pep from pyruvate (pyr), 1 ATP is required, and converting 3-phospho-glycerate (3pg) into 3-phospho-glyceroyl phosphate (13dpg) requires another ATP. Similarly, making glyceraldehyde 3-phosphate (g3p) from 13dpg consumes one NADH. Moreover, one ATP is required to activate acetate to acetyl-CoA (accoa). The *i*JO1366 model predicted that most energy molecules would be produced from the TCA cycle, and the PP pathway would be barely used in energy production on acetate or other poor carbon sources. The number of energy molecules that were calculated to be generated from the TCA cycle is sufficient to accommodate the energy requirements for gluconeogenesis and acetate conversion. However, NADH or NADPH needs to be converted into ATP, since the major energy expenditure would occur with ATP. Following the fluxes coming from the TCA cycle, the model-based simulation sheds light on how NADH could be converted into ATP when cells are growing on the poor carbon sources. NADH oxidoreductase I uses NADH to pump out proton molecules into the periplasm so that ATP synthase can generate ATP from the proton gradient (Figure 5E).

Since the i*J*O1366 model predicted there would be an increased flux though NADH oxidoreductase I, it was postulated that Cra or CRP may be involved in regulating the expression of the enzyme complex. NADH oxidoreductase I is encoded by a long operon, *nuoABCEFGHIJKLMN,* and ChIP-exo experiments provided evidence that Cra binds upstream of this operon, indicating regulation by Cra (Figure 5F). To determine if this regulation is positive or negative, expression change of this operon upon *cra* knock-out was analyzed (Figure 5G). On glucose, knocking out *cra* did not change the expression of *nuoABCEFGHIJKLMN,* however the expression significantly decreased on acetate (ranksum test p-value < 1.5X10^−5^). Thus, Cra up-regulates the expression of *nuoABCEFGHIJKLMN.*

Cra was reported to regulate a broad range of metabolic genes, but independent of CRP (12). However, there is supporting evidence that Cra directly regulates the expression of *crp* (16,37). The Cra ChIP-exo dataset supports its binding upstream of σ^70^-dependent promoter of *crp* (Figure S8), however the expression change of *crp* was not significant between glucose and acetate (Figure S6) nor between WT and *Δcra* (Figure S11). No evidence has been found that CRP regulates the expression of *cra.* Thus, the regulatory interaction between CRP and Cra is responsible for the competition between them on the expression of target genes that they both regulate.

## DISCUSSION

In this study, the complex transcriptional regulatory network of carbon metabolism in enterobacteria was investigated using a combination of genome-wide experimental measurements and computer simulation of a genome-scale metabolic model. The ChIP-exo and RNA-seq methods were applied to Cra when *E. coli* was grown on glucose, fructose, and acetate, and led to the identification of 97 genes in the Cra regulon. The definition of the Cra regulon showed that Cra and CRP have distinct roles in carbon metabolism regulation. Cra is involved in the repression of glycolysis, while historical data shows that CRP is focused on the activation of the TCA cycle. Expression profiling illustrated that the expression of genes in glycolysis is highest on fructose, and genes in the TCA cycle were more highly expressed on acetate. Model-based simulation and flux balance analysis were employed to explain this observation, and it was found that it is due to the fact that energy production is mostly coming from the TCA cycle. This energy production from the TCA cycle enables gluconeogenesis when growing on unfavorable carbon sources. The conversion of energy molecules from NADH to ATP happens during this process, and this explains Cra regulation on the redox pathway. A single base-pair resolution of the experimental methods and detailed sequence analysis on Cra and CRP binding sites clarified how the activity of Cra overrides the activity of CRP in regulation of their target genes. The optimal gene expression on different carbon sources could be implemented by differential activation of Cra and CRP on glucose and fructose, and Cra activity overriding CRP binding on unfavorable carbon sources. Conservation analysis demonstrated that transcriptional regulation by Cra might be a more widely used strategy in modulating carbon and energy metabolism over regulation by CRP in microorganisms.

Most Cra regulon genes are metabolic enzymes; however there are three TFs in its regulon: *pdhR, nikR,* and *baeR.* While the affiliation of *nikR* or *baeR* to carbon metabolism is still unclear, *pdhR* is involved in carbon metabolism by sensing pyruvate (38). The binding sites of PdhR have been investigated with both *in vitro* (39) and *in vivo* (40) methods. In both studies, *ndh* was annotated as a PdhR target gene. *ndh* encodes NADH oxidoreductase II (NDH-2) which is one of two distinct NADH dehydrogenases in *E. coli.* The other NADH dehydrogenase is NDH-1 that is encoded by *nuo* genes, and Cra regulates the expression of the *nuo* operon. Moreover, Cra and PdhR both regulate *cyoABCDE,* which encodes cytochrome *bo* oxidase. How the involvement of the electron transport system is relevant to growth on pyruvate has not been fully elaborated. However, it makes sense that the optimal growth on unfavorable carbon sources accompanies regulation on the redox pathway. It is possible that this may be because of energy production from the TCA cycle and conversion between energy molecules as similarly shown on acetate for Cra.

In summary, cutting-edge experimental measurements with ChIP-exo and RNA-seq provided the regulatory information for Cra on the core carbon metabolism at the genome-scale. Integration of this experimentally-derived regulatory information and *in silico* flux calculation with a genome-scale metabolic model expanded the scope of carbon metabolism regulation by Cra. Cra supports the optimal cell growth on the poor carbon sources by at least three mechanisms (Figure 6). First, Cra redirects the enzymatic flux through glycolysis towards gluconeogenesis, but more importantly it decreases the flux volume through this pathway. Second, Cra activates the activity of the glyoxylate shunt pathway together with activation of the TCA cycle. Third, Cra up-regulates some components in the respiratory chain to provide the energy balance between the repressed glycolysis pathway and the activated TCA cycle. Most importantly, the repression of the glycolysis pathway and the activation of glyoxylate shunt pathway crucially depend on the overriding regulatory activity of Cra over that of CRP.

**Figure 6.**
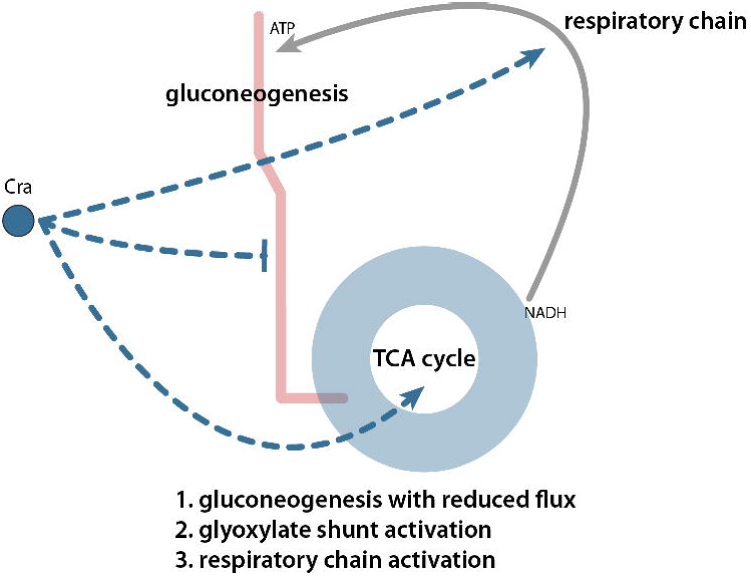
Expanded regulatory roles of Cra with its overriding effect on CRP regulation. Cra redirects the flux of the glycolysis pathway towards gluconeogenesis with a reduced amount of flux, and increases the reaction flux of the glyoxylate shunt pathway. For those pathways, the overriding regulatory effect of Cra over CRP ensures the optimal growth of *E. coli.* In addition, the previously unknown regulation by Cra on the components of the respiratory chain enables converting between energy molecules, balancing increased flux through TCA cycle and decrease flux through gluconeogenesis.

The consolidation of the experimental measurement of *in vivo* states of transcriptional components and the computational prediction of *in silico* states of metabolic activities makes for an integrated genome-scale approach with which to investigate the network level mechanisms of transcriptional regulation in bacteria. Experimental measurements with recently developed methods at the single base-pair resolution enable researchers to determine the transcriptional regulation activity and to follow biological questions from the dataset. However, experimental methods can only provide a monolithic snapshot of internal *in vivo* states of transcriptional regulation under the given conditions. Model-based *in silico* simulation, on the other hand, allows researchers to investigate the activity of a reaction in association with other connected reactions and to explore feasible cellular states. Thus it is possible to put biological questions or findings in a broader or expanded context. For instance, the linkage found in this study could be further investigated in the context of carbon and redox metabolism (41) in combinatorial conditions, which would contribute to understanding carbon metabolism regulation in the context of oxygen-limiting conditions. Thus, elucidation of transcriptional regulation of the core carbon metabolism in bacteria exhibited the benefits from combining genome-wide experimental measurement and simulation with a genome-scale metabolic model.

## AUTHOR CONTRIBUTIONS

DK, BOP conceived the study. SWS, DK, GIG, YG performed experiments. DK and HN performed the computational analysis. BOP supervised the study. DK, SWS, HN, YG, and BOP wrote the manuscript. All authors helped edit the final manuscript.

## ACKNOWLEDGEMENTS

We thank Marc Abrams for helpful assistance in writing and editing the manuscript. This research was supported by the grant NNF16CC0021858 from Novo Nordisk Foundation Center for Biosustainability at the Danish Technical University and by NIH NIGMS (National Institute of General Medical Sciences) grant GM102098. This research used resources of the National Energy Research Scientific Computing Center, which is supported by the Office of Science of the US Department of Energy under Contract No. DE-AC02-05CH11231. The whole dataset of ChIP-exo and RNA-seq has been deposited to GEO with the accession number of GSE65643.

## CONFLICT OF INTEREST

The authors declare no conflict of interest.

## STAR Methods

All strains used in this study are *E. coli* K-12 MG1655 and its derivatives. For ChIP-exo experiments, the *E. coli* strain harboring cra-8myc was generated as described previously (42), and was grown on glucose, fructose, and acetate to perform ChIP-exo as previously described (17). For RNA-seq experiments, a deletion mutant Δ*cra* was constructed by λ red-mediated site-specific recombination system (43). The wild type and Δ*cra* were grown on glucose, fructose, and acetate to perform RNA-seq as previously described (17). Calculation of differentially expressed genes was conducted by using bowtie (44) and Cuffdiff (45). ChIP-exo reads were processed with MACE software (46) (https://code.google.com/p/chip-exo/). Flux analysis to calculate fluxes through metabolic reactions under different nitrogen sources was performed with *E. coli* M model (20) and COBRApy (47). More detailed procedures are described in Supplementary Experimental Procedures section.

## SUPPLEMENTAL INFORMATION

### Supplementary Experimental Procedures

### Bacterial strains, media, and growth conditions

All strains used in this study are *E. coli* K-12 MG1655 and its derivatives. For ChIP-exo experiments, the *E. coli* strain harboring cra-8myc was generated as described previously (42). For growth rate measurement and expression profiling by RNA-seq, deletion mutant Δ*cra* and Δ*crp* were constructed by λ red-mediated site-specific recombination system (43). For growth rate measurement, glycerol stocks of *E. coli* strains were inoculated into M9 minimal media with different carbon sources, glucose, fructose, galactose, succinate, glycerol or acetate. The concentration of carbon sources was 0.2% (w/v). M9 minimal media was also supplemented with 1 ml trace element solution (100X) containing 1 g EDTA, 29 mg ZnSO_4_.7H_2_O, 198 mg MnCl_2_.4H_2_O, 254 mg CoCl_2_.6H_2_O, 13.4 mg CuCl_2_, and 147 mg CaCl_2_. The culture was incubated at 37°C overnight with agitation, and then was used to inoculate the fresh media. For RNA-seq expression profiling, glycerol stocks of *E. coli* strains were inoculated into M9 minimal media with different carbon sources, glucose, fructose or acetate. The concentration of carbon sources was 0.2% (w/v). M9 minimal media was also supplemented with 1 ml trace element solution (100X). The culture was incubated at 37°C overnight with agitation, and then was used to inoculate the fresh media. The fresh culture was incubated at 37°C with agitation to the mid-log phase (OD_600_≈ 0.5 for glucose and fructose, and OD_600_≈ 0.25 for acetate).

### ChIP-exo experiment

ChIP-exo experiment was performed following the procedures previously described (17). In brief, to identify Cra binding maps *in vivo,* we isolated the DNA bound to Cra from formaldehyde cross-linked *E. coli* cells by chromatin immunoprecipitation (ChIP) with the specific antibodies that specifically recognizes myc tag (9E10, Santa Cruz Biotechnology), and Dynabeads Pan Mouse IgG magnetic beads (Invitrogen) followed by stringent washings as described previously (27). ChIP materials (chromatin-beads) were used to perform on-bead enzymatic reactions of the ChIP-exo method (17,48). Briefly, the sheared DNA of chromatin-beads was repaired by the NEBNext End Repair Module (New England Biolabs) followed by the addition of a single dA overhang and ligation of the first adaptor (5’-phosphorylated) using dA-Tailing Module (New England Biolabs) and NEBNext Quick Ligation Module (New England Biolabs), respectively. Nick repair was performed by using PreCR Repair Mix (New England Biolabs). Lambda exonuclease- and RecJf exonuclease-treated chromatin was eluted from the beads and the protein-DNA cross-link was reversed by overnight incubation at 65°C. RNAs- and Proteins-removed DNA samples were used to perform primer extension and second adaptor ligation with following modifications. The DNA samples incubated for primer extension as described previously (17) were treated with dA-Tailing Module (New England Biolabs) and NEBNext Quick Ligation Module (New England Biolabs) for second adaptor ligation. The DNA sample purified by GeneRead Size Selection Kit (Qiagen) was enriched by polymerase chain reaction (PCR) using Phusion High-Fidelity DNA Polymerase (New England Biolabs). The amplified DNA samples were purified again by GeneRead Size Selection Kit (Qiagen) and quantified using Qubit dsDNA HS Assay Kit (Life Technologies). Quality of the DNA sample was checked by running Agilent High Sensitivity DNA Kit using Agilent 2100 Bioanalyzer (Agilent) before sequenced using MiSeq (Illumina) in accordance with the manufacturer’s instructions. Each modified step was also performed in accordance with the manufacturer’s instructions. ChIP-exo experiments were performed in biological duplicate.

### RNA-seq expression profiling

Three milliliters of cells from mid-log phase culture were mixed with 6 ml RNAprotect Bacteria Reagent (Qiagen). Samples were mixed immediately by vortexing for 5 seconds, incubated for 5 minutes at room temperature, and then centrifuged at 5000×g for 10 minutes. The supernatant was decanted and any residual supernatant was removed by inverting the tube once onto a paper towel. Total RNA samples were then isolated using RNeasy Plus Mini kit (Qiagen) in accordance with the manufacturer’s instruction. Samples were then quantified using a NanoDrop 1000 spectrophotometer (Thermo Scientific) and quality of the isolated RNA was checked by running RNA 6000 Pico Kit using Agilent 2100 Bioanalyzer (Agilent).

Paired-end, strand-specific RNA-seq was performed using the dUTP method (49) with the following modifications which is previously described (17). The ribosomal RNAs were removed from 2 µg of isolated total RNA with Ribo-Zero rRNA Removal Kit (Epicentre) in accordance with the manufacturer’s instruction. Subtracted RNA was fragmented for 2.5 min at 70 °C with RNA Fragmentation Reagents (Ambion), and then fragmented RNA was recovered with ethanol precipitation. Random primer (3 µg) and fragmented RNA in 4 µl was incubated in 5 µl total volume at 70 °C for 10 minutes, and cDNA or the first strand was synthesized using SuperScript III first-strand synthesis protocol (Invitrogen). The cDNA was recovered by phenol-chloroform extraction followed by ethanol precipitation. The second strand was synthesized from this cDNA with 20 μl of fragmented cDNA:RNA, 4 μl of 5× first strand buffer, 30 μl of 5× second strand buffer, 4 μl of 10 mM dNTP with dUTP instead of dTTP, 2 μl of 100 mM DTT, 4 μl of *E. coli* DNA polymerase (Invitrogen), 1 μl of *E. coli* DNA ligase (Invitrogen), 1 μl of *E. coli* RNase H (Invitrogen) in 150 μl of total volume. This reaction mixture was incubated at 16 °C for 2 hours, and fragmented DNA was recovered with PCR clean-up kit (QIAGEN) and eluted in 30 μl of nuclease-free water. The fragmented DNA was end-repaired with End Repair Kit (New England Biolabs), and dA-tailed with dA-Tailing Kit (New England Biolabs), and then ligated with 7.5 μg of DNA adaptor mixture with Quick Ligation Kit (New England Biolabs). The adaptor-ligated DNA was size-selected to removed un-ligated adaptors with GeneRead Size Selection Kit (QIAGEN), and treated with 1 U of USER enzyme (New England Biolabs) in 30 μl of total volume, and incubated at 37 °C for 15 minutes followed by 5 minutes at 95 °C. The USER-treated DNA was amplified by PCR to generate sequencing library for Illumina sequencing. The samples were sequenced using MiSeq (Illumina) in accordance with the manufacturer’s instructions. All RNA-seq experiments were performed in biological duplicate.

### Peak calling for ChIP-exo dataset

Peak calling was performed as previously described(17). Sequence reads generated from ChIP-exo were mapped onto the reference genome (NC_000913.2) using bowtie(44) with default options to generate SAM output files (Table S6). MACE program(46) was used to define peak candidates from biological duplicates for each experimental condition with sequence depth normalization. To reduce false-positive peaks, peaks with signal-to-noise (S/N) ratio less than 1.5 were removed. The noise level was set to the top 5% of signals at genomic positions because top 5% makes a background level in plateau and top 5% intensities from each ChIP-exo replicates across conditions correlate well with the total number of reads(17–19). The calculation of S/N ratio resembles the way to calculate ChIP-chip peak intensity where IP signal was divided by Mock signal. Then, each peak was assigned to the nearest gene. Genome-scale data were visualized using MetaScope (http://systemsbiology.ucsd.edu/Downloads/MetaScope).

### Motif search from ChIP-exo peaks

The sequence motif analysis for TFs and σ-factors was performed using the MEME software suite (50). For Cra, sequences in binding regions were extracted from the reference sequence (NC_000913.2).

### Calculation of differentially expressed gene

Sequence reads generated from RNA-seq were mapped onto the reference genome (NC_000913.2) using bowtie (44) with the maximum insert size of 1000 bp, and 2 maximum mismatches after trimming 3 bp at 3’ ends (Table S7). SAM files generated from bowtie, then, were then used for Cufflinks (http://cufflinks.cbcb.umd.edu/) (45) to calculate fragments per kilobase of exon per million fragments (FPKM). Cufflinks was run with default options with the library type of dUTP RNA-seq and the default normalization method (classic-fpkm). Differentially expressed genes were calculated with DESeq2(51) and expression with log2 fold change ≥ 1.0 and adjusted p-value ≤ 0.05 was considered as differentially expressed. Genome-scale data were visualized using MetaScope (http://svstemsbiology.ucsd.edu/Downloads/MetaScope).

### COG functional enrichment

Cra regulons were categorized according to their annotated clusters of orthologous groups (COG) category. Functional enrichment of COG categories in Cra target genes was determined by performing hypergeometric test, and *p-value*< 0.05 was considered significant.

### FBA analysis and MCMC sampling to calculate the metabolic flux

FBA analysis and MCMC sampling was performed with iJO1366 *E. coli* metabolic model (20), COBRA Toolbox v2.0 (52) and COBRApy (47) as previously described (53). In brief, the distribution of feasible fluxes for each reaction in the iJO1366 model was determined using Markov chain Monte Carlo (MCMC) sampling (54). Specifically, uptake rates for the carbon sources were measured with HPLC and were used to constrain the model: -8.437 mmol/gDW/hr for glucose, -7.546 mmol/gDW/hr for fructose, and -7.671 mmol/gDW/hr for acetate. The biomass objective function (a proxy for growth rate) was provided a lower bound of 95% of the optimal growth rate as computed by FBA. Thus, the sample flux distributions by MCMC sampling method represented sub-optimal flux distributions. MCMC sampling was used to obtain 10 thousands of feasible flux distributions, and the average of flux samples for each reaction was used. Sampled points of reactions in loops were removed before further analysis. Reactions in loops were calculated by using flux variability analysis (FVA) on iJO1366 model.

### Conservation analysis of Cra and CRP regulon genes

Gene annotation of 552 species and strains ranging from *Escherichia* to archaea, were obtained from the SEED server (http://theseed.org) and ortholog calculation to *E. coli* K-12 MG1655 was performed on RAST (Rapid Annotation using Subsystem Technology) server(55). From RAST output, orthologous genes with bi-directional hits were only retained. Conservation level of Cra and CRP regulon genes in carbon metabolism were calculated from orthologs retained from RAST output.

### Supplementary Tables

**Table S1.** Cra binding sites

**Table S2.** Expression profiling of WT and Δcra on three carbon sources

**Table S3.** Expression change of Cra regulon

**Table S4.** Regulatory mode on Tus

**Table S5.** Relative Expression of Genes in Glycolysis and TCA cycle

**Table S6.** Statistics on ChIP-exo sequencing result

**Table S7.** Statistics on RNA-seq sequencing result

### Supplementary Figures

**Figure S1.**
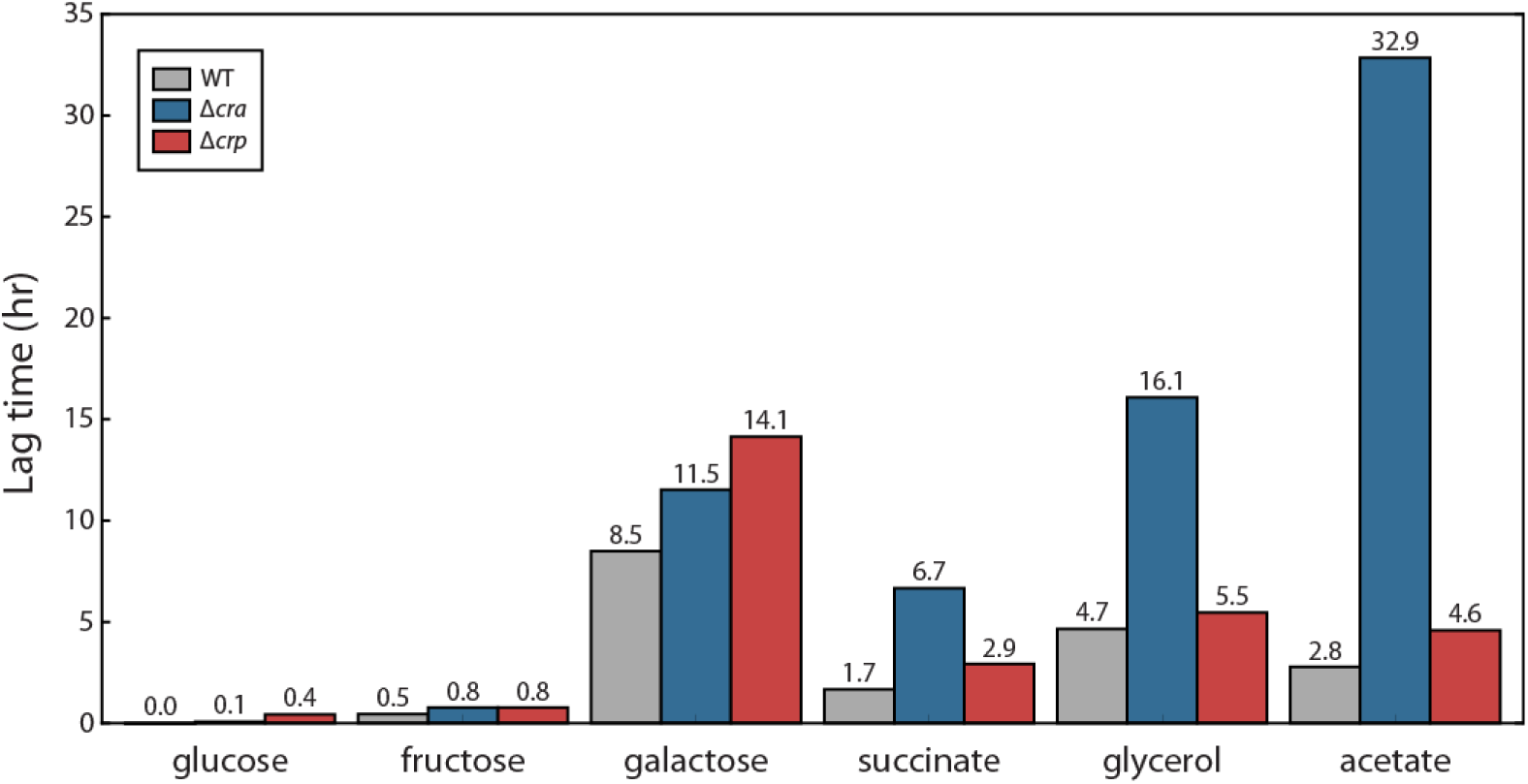
Measurement of lag-phase time of *E. coli* WT, Δ*cra,* and Δ*crp* on 6 different carbon sources. Disruption of *cra* showed a much longer lag-phase time on less favorable carbon sources, succinate, glycerol, and acetate, than that of *crp.*

**Figure S2.**
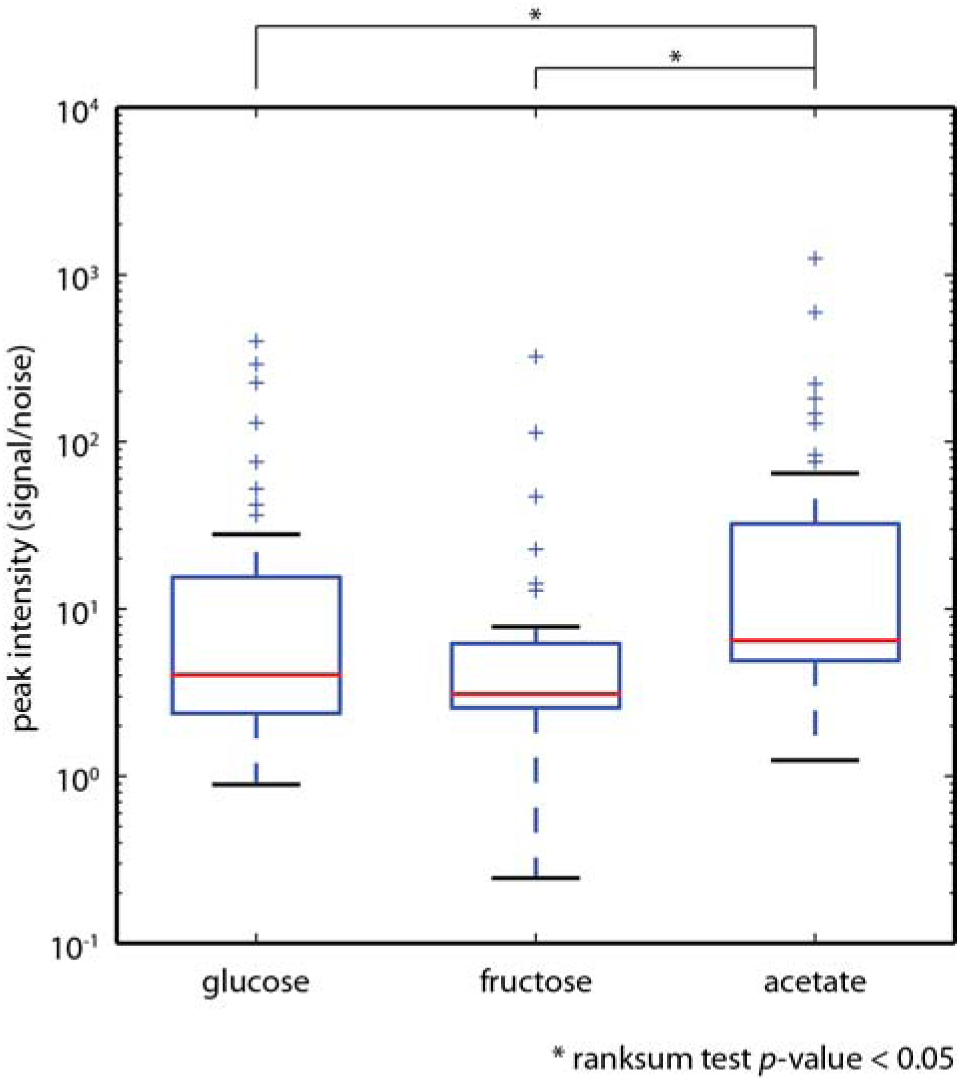
Peak intensity of Cra ChIP-exo binding sites on glucose, fructose, and acetate. Among 3 carbon sources, peak intensity was strongest on acetate, and was weakest on fructose. Differences of peak intensities between glucose/acetate and fructose/acetate were statistically significant (* indicates ranksum test p-value < 0.05).

**Figure S3.**
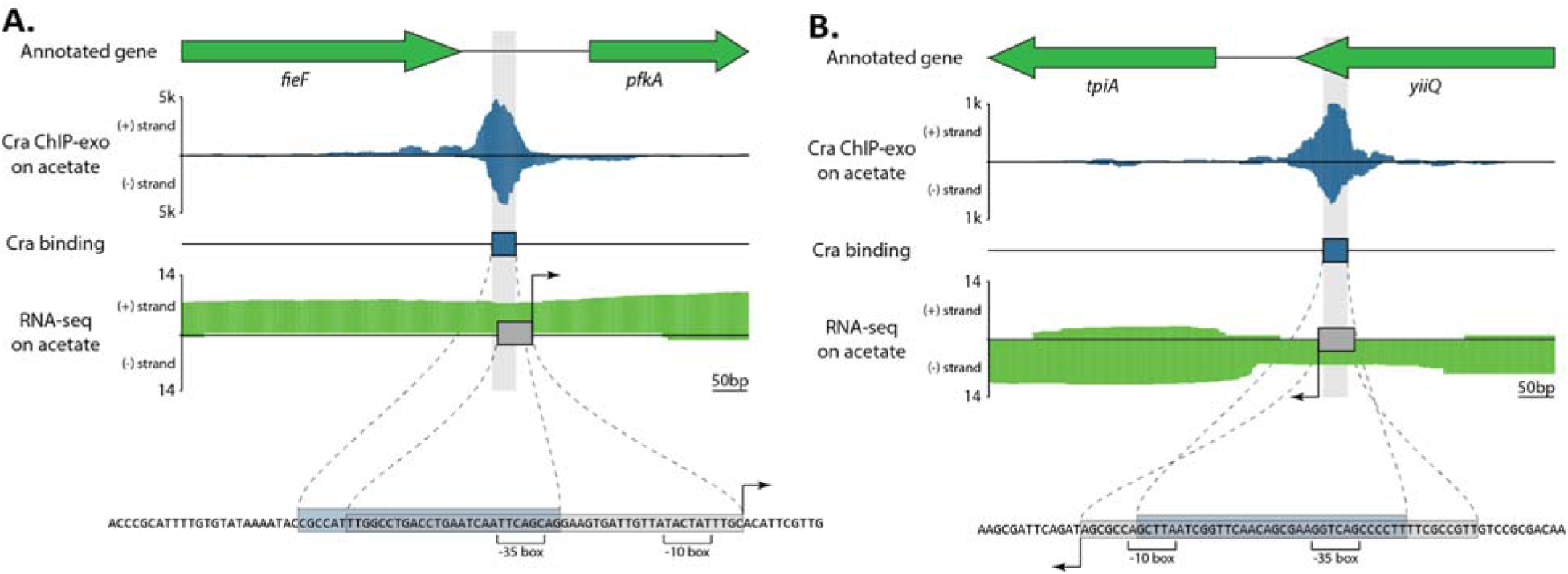
Zoomed-in examples of Cra binding sites upstream of *pfkA* and *tpiA.* ChIP-exo experiments provide a better resolution over long-established ChIP methods, such as ChIP-chip or ChIP-seq. Two Cra binding sites identified by ChIP-exo are overlapping with promoters, indicating possible repression on expression of *pfkA* and *tpiA.*

**Figure S4.**
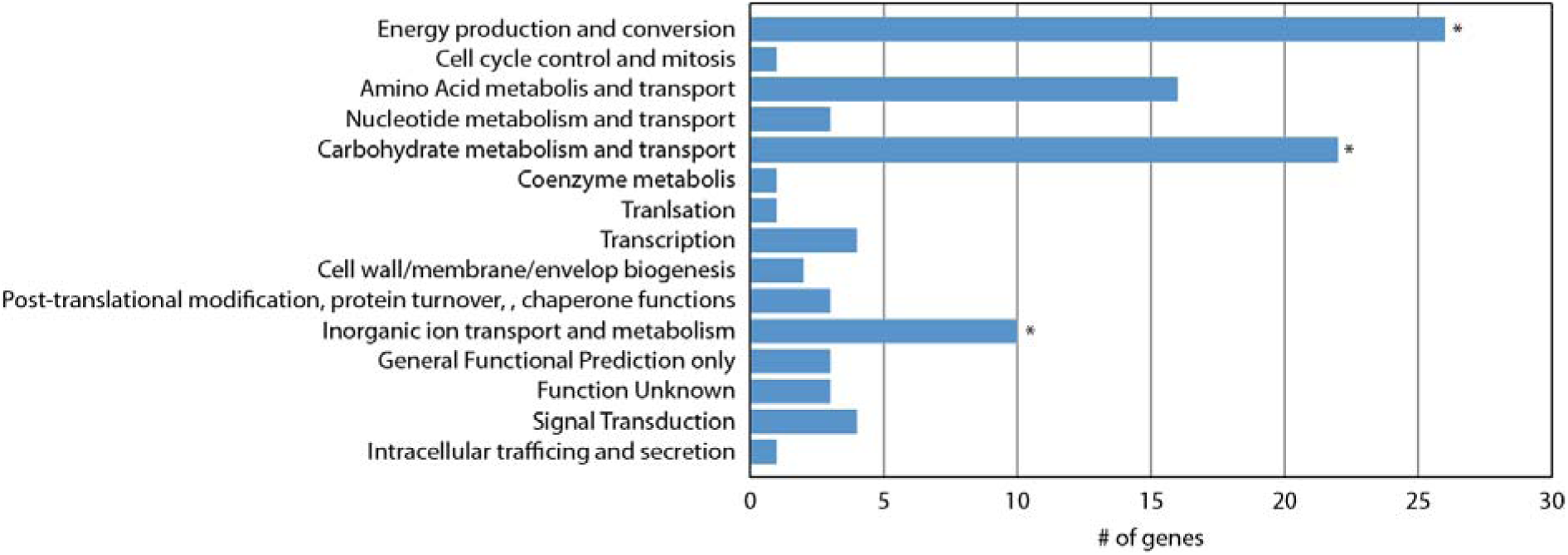
COG analysis of Cra regulon genes. Cra regulon has enriched functions in 3 groups, energy production/conversion, carbohydrate metabolism/transport, and inorganic ion transport/metabolism (* indicates hypergeometric test p-value < 0.05). Hypergeometric test p-values for “Energy production and conversion” and “Carbohydrate metabolism and transport” were < 10^−6^, and *p*-value for “Inorganic ion transport and metabolism” was 3.6X10^−2^.

**Figure S5.**
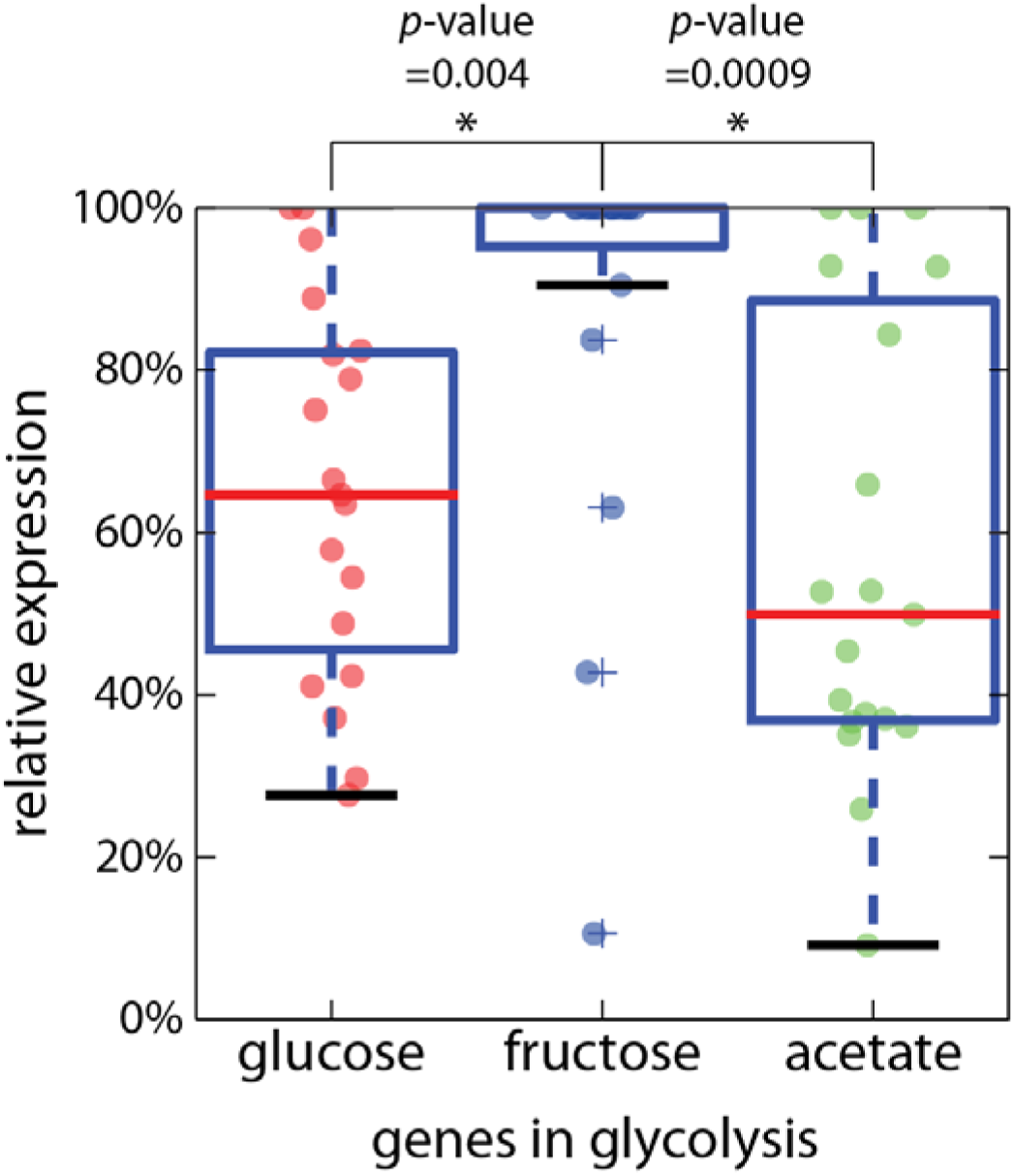
Comparison of the relative expression of all genes, glycolytic and gluconeogenic, in glycolysis. Even if genes in gluconeogenic genes were included, the same pattern shows up in comparison to the relative expression of glycolysis genes between glucose, fructose, and acetate. On average, genes in glycolysis are more expressed on fructose, and are the least expressed on acetate. These comparisons were all statistically significant (* indicates ranksum p-value < 0.05).

**Figure S6.**
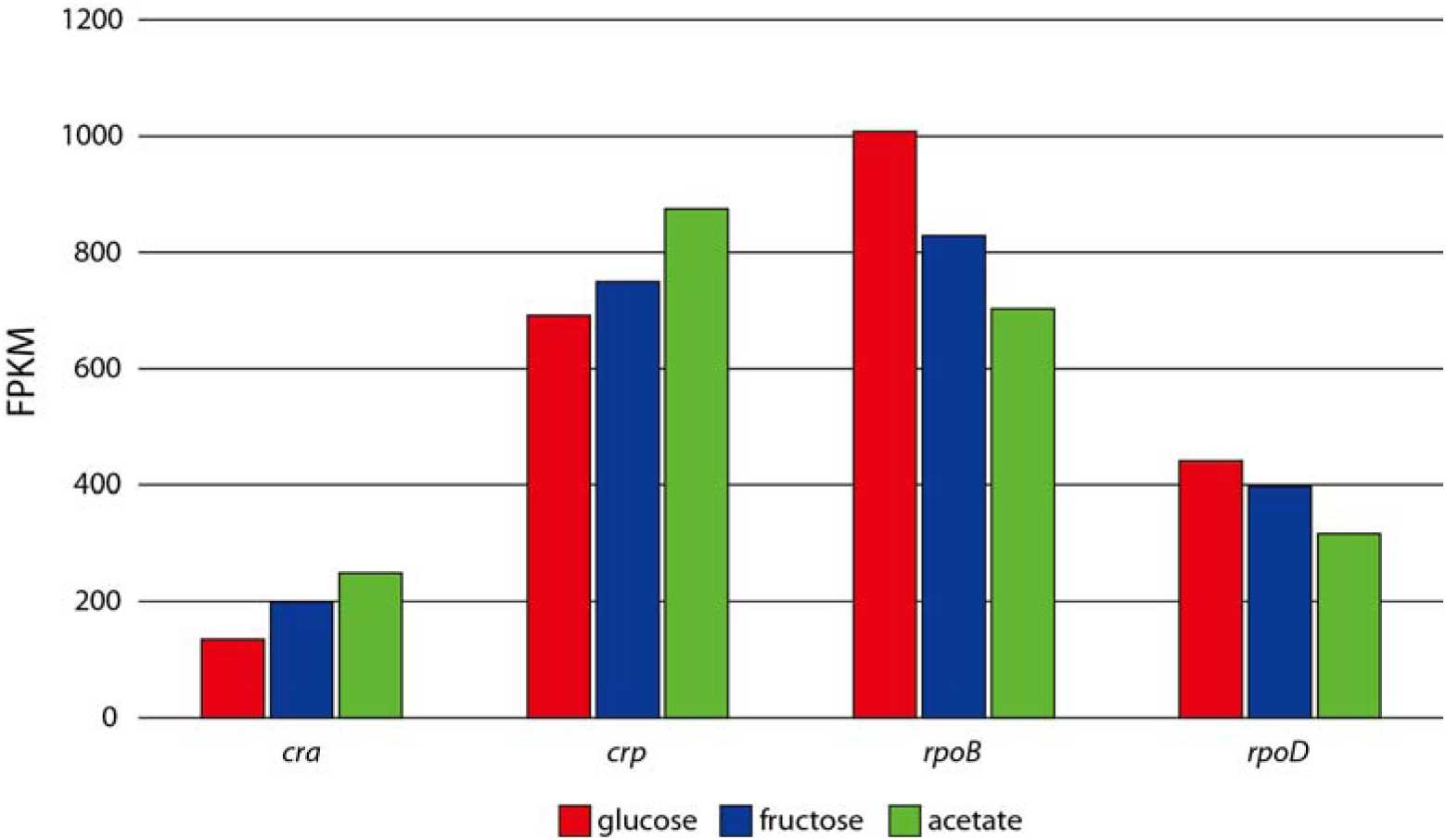
Expression comparison of *cra* and *crp* on different carbon sources. Expression change of *cra* and *crp* did not change significantly (fold-change ≥2). *rpoB* and *rpoD,* subunits of RNA polymerase, were presented as controls, since their expression was not supposed to change significantly.

**Figure S7.**
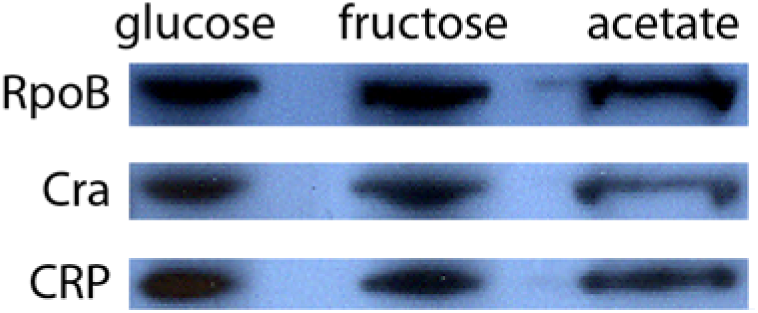
Protein expression comparison of *cra* and *crp* on different carbon sources. Protein expression level of Cra and CRP was compared by western blotting. RpoB was used as a control, which is expected to show constant level of protein expression.

**Figure S8.**
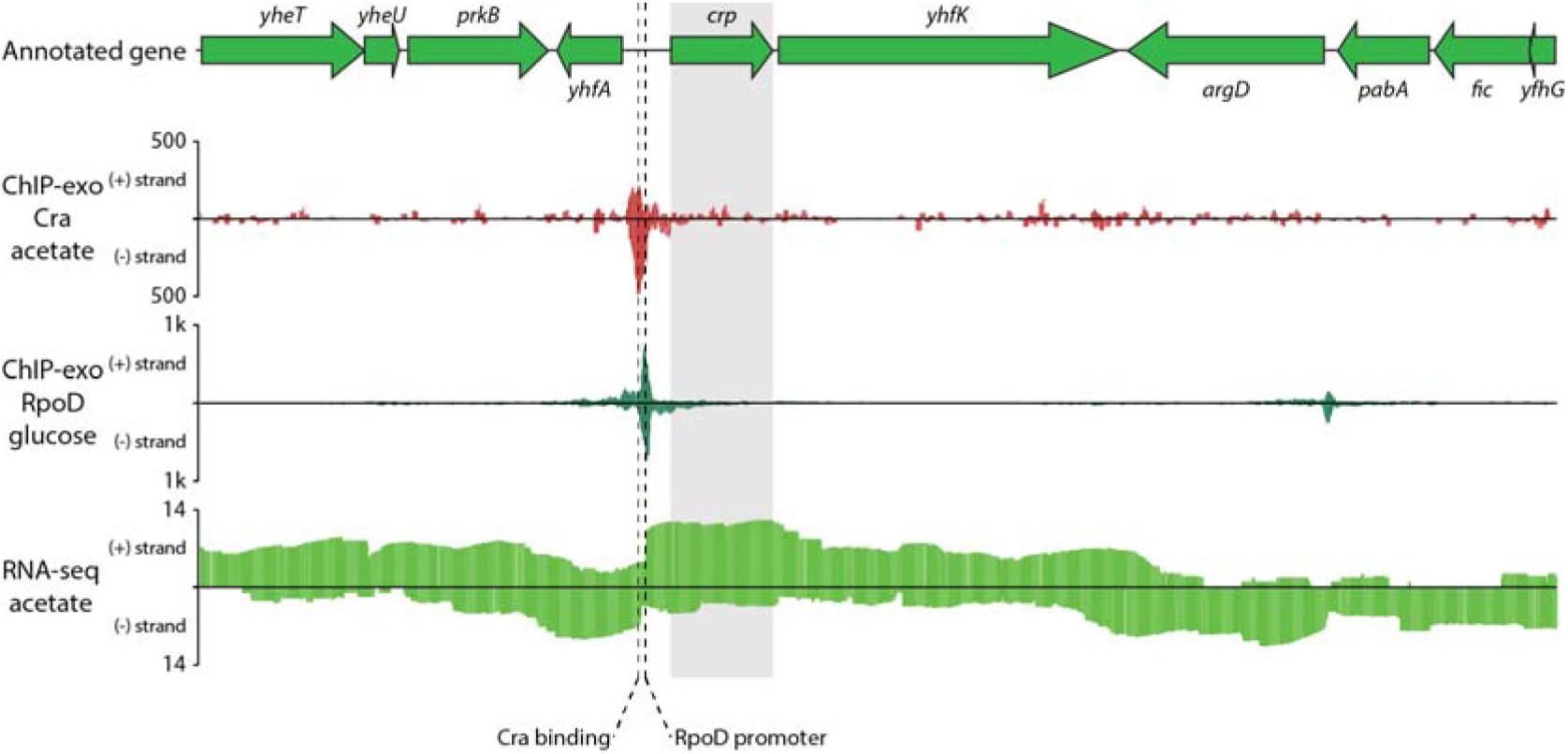
ChIP-exo binding of Cra and expression profiling in the region surrounding *crp.* Cra binding upstream of *crp* was observed from Cra ChIP-exo dataset. RpoD ChIP-exo dataset (unpublished) presents the presence of σ^70^-dependent promoter slightly downstream of the Cra binding site.

**Figure S9.**
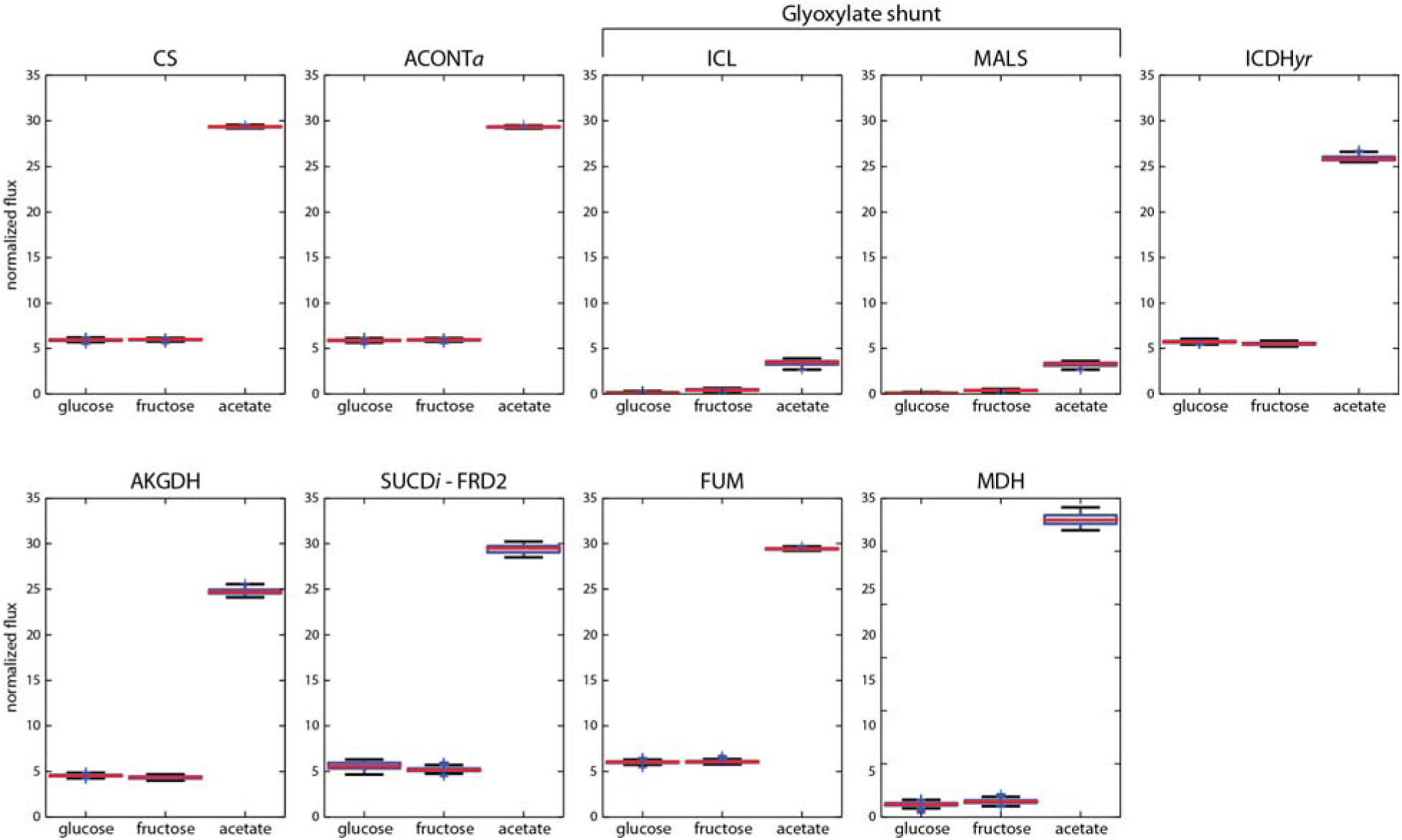
Simulated fluxes though reactions in the TCA cycle. In every reaction in the TCA cycle, fluxes were activated on acetate when compared to glucose or fructose.

**Figure S10.**
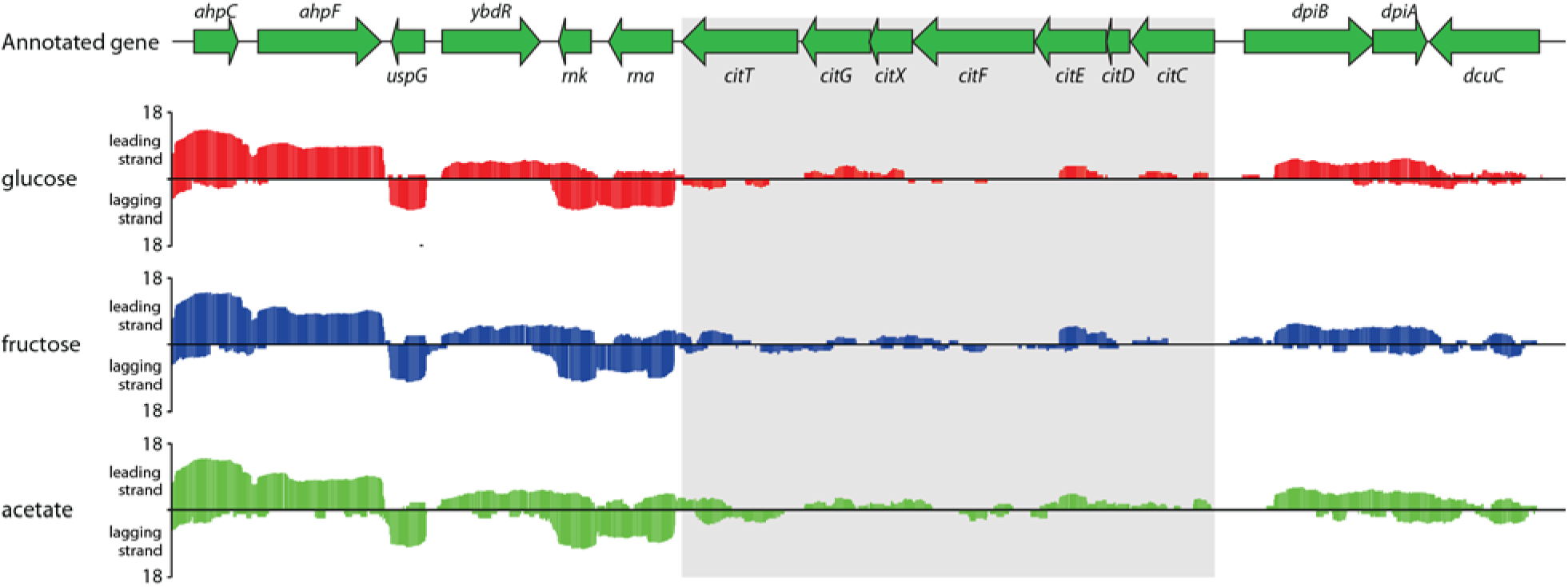
Expression profiling of *citCDEFXGT* operon on different carbon sources. The operon *citCDEFXGT* was barely expressed on glucose, fructose, and acetate.

**Figure S11.**
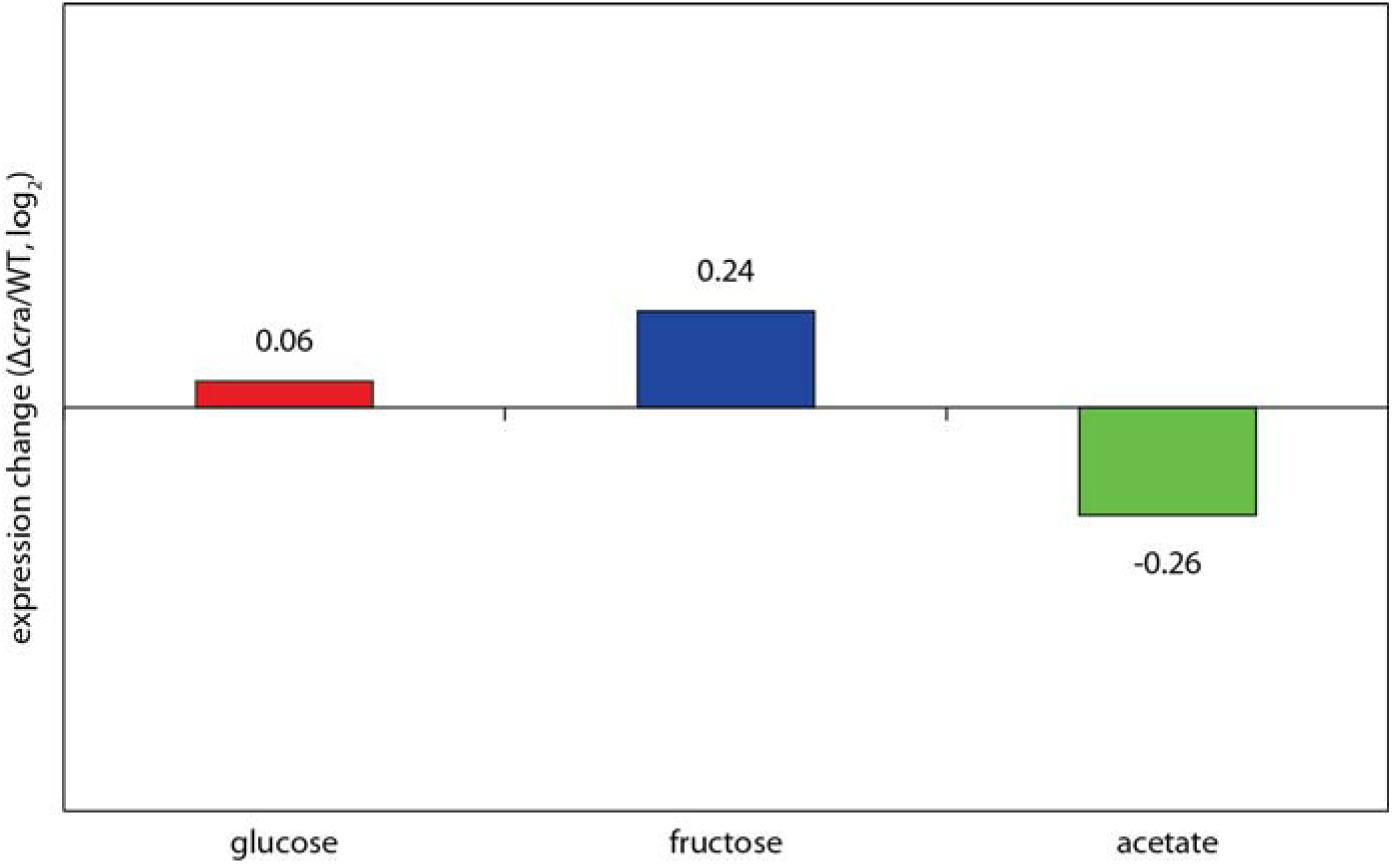
Expression change of *crp* in *E. coli* WT and Δ*cra* on glucose, fructose, and acetate. Knocking out *cra* did not change the expression of *crp* significantly on all carbon sources.

